# Novel truncating mutations in *CTNND1* cause a dominant craniofacial and cardiac syndrome

**DOI:** 10.1101/711184

**Authors:** Reham Alharatani, Athina Ververi, Ana Beleza-Meireles, Weizhen Ji, Emily Mis, Quinten T. Patterson, John N. Griffin, Nabina Bhujel, Caitlin A. Chang, Abhijit Dixit, Monica Konstantino, Christopher Healy, Sumayyah Hannan, Natsuko Neo, Alex Cash, Dong Li, Elizabeth Bhoj, Elaine H. Zackai, Ruth Cleaver, Diana Baralle, Meriel McEntagart, Ruth Newbury-Ecob, Richard Scott, Jane A. Hurst, Ping Yee Billie Au, Marie Therese Hosey, Mustafa Khokha, Denise K. Marciano, Saquib A. Lakhani, Karen J. Liu

**Affiliations:** Centre for Craniofacial and Regenerative Biology, King’s College London, UK; Paediatric Dentistry, Centre of Oral, Clinical and Translational Science, King’s College London, UK; Clinical Genetics, Great Ormond Street Hospital, London, UK; Genetics Department, Guy’s and St. Thomas’ NHS Foundation Trust, London, UK; Pediatric Genomics Discovery Program, Department of Pediatrics, Yale University School of Medicine, New Haven, CT 06520, USA; Departments of Medicine and Cell Biology, University of Texas Southwestern Medical Center, Texas, USA; Pediatric Genomics Discovery Program, Departments of Genetics and Pediatrics, Yale University School of Medicine, New Haven, CT 06520, USA; South Thames Cleft Unit, Guy’s and St. Thomas’ NHS Foundation Trust, London, UK; University of Calgary, Alberta Children’s Hospital Research Institute, Calgary, Alberta, Canada; Nottingham University Hospitals NHS Trust, City Hospital Campus, Nottingham, UK; Tokyo Medical and Dental University, Tokyo, Japan; Department of Pediatrics, Division of Human Genetics, Children’s Hospital of Philadelphia, USA; Peninsula Clinical Genetics Service, Royal Devon and Exeter NHS Foundation Trust, Exeter, UK; Human Development and Health, Faculty of Medicine, University of Southampton, Southampton, United Kingdom; Department of Clinical Genetics, St George’s Hospital, London, UK; Clinical Genetics, University Hospital Bristol NHS Foundation Trust, Bristol, UK

**Keywords:** *CTNND1*, p120-catenin, craniofacial, cardiac, blepharocheilodontic syndrome, cleft lip and palate, hypodontia, neural, larynx, *Xenopus;* mouse

## Abstract

*CTNND1* encodes the p120-catenin (p120) protein, which has a wide range of functions, including the maintenance of cell-cell junctions, regulation of the epithelial-mesenchymal transition and transcriptional signaling. Due to advances in next generation sequencing, *CTNND1* has been implicated in human diseases including cleft palate and blepharocheilodontic syndrome (BCD) albeit only recently. In this study, we identify eight novel protein-truncating variants, six *de novo,* in thirteen participants presenting with craniofacial dysmorphisms including cleft palate and hypodontia, as well as congenital cardiac anomalies, limb dysmorphologies and neurodevelopmental disorders. Using conditional deletions in mice as well as CRISPR/Cas9 approaches to target *CTNND1* in *Xenopus*, we identified a subset of phenotypes that can be linked to p120-catenin in epithelial integrity and turnover, and additional phenotypes that suggest mesenchymal roles of *CTNND1.* We propose that *CTNND1* variants have a wider developmental role than previously described, and that variations in this gene underlie not only cleft palate and BCD but may be expanded to a broader velocardiofacial-like syndrome.

## Introduction

Genetic variation in *CTNND1,* which encodes for the armadillo-repeat protein p120-catenin (p120), is associated with human birth defects, most notably non-syndromic cleft palate and blepharocheilodontic (BCD) syndrome, which involves eyelid, lip and tooth anomalies [MIM: 617681]^1–3^. In contrast, *CTNND1* has broader developmental roles in animal models. For example, conditional deletions in mice demonstrate the importance of *CTNND1* for development not only for skin and teeth, but also for kidneys and other structures^4–10^, and complete deletion of *CTNND1* leads to prenatal lethality^5, 9^. Similarly, loss-of-function experiments in *Xenopus* implicate *CTNND1* in craniofacial development^11, 12^. Here, we describe a series of patients with *CTNND1* variants, all of whom present with multisystem involvement that demonstrates a broad spectrum craniofacial and cardiac syndrome.

p120-catenin is a member of the catenin superfamily of proteins studied in catenin-cadherin interactions; notably, it binds to and stabilizes E-cadherin (*CDH1*) at junctional complexes in epithelia^13–17^. This binding is via the p120-catenin armadillo repeat domain, and displacement of p120-catenin from E-cadherin is a key regulatory event at the adherens junction, that results in endocytosis of E-cadherin and loss of the junction. The protein has a second function as a scaffolding protein for the GTPase RhoA and associated Rho regulatory proteins^18, 19^. In addition, it can also directly interact with the zinc finger transcriptional repressor Kaiso (ZBTB33), facilitating Wnt signal transduction^20, 21^. Thus, p120-catenin appears to be a multi-functional protein, promoting epithelial stability when in complex with E-cadherin, and regulating RhoA and transcriptional activities. p120-catenin is also able to associate with mesenchymal cadherins such as N-cadherin and cadherin-11^17, 22^. In mesenchymal cells, p120-catenin associates with non-epithelial cadherins, regulating motility and invasion via cytoskeletal events and transcription. Given its functions in both epithelia and mesenchyme, it is unsurprising that both loss and gain of p120-catenin have been associated with oncogenesis^23–25^.

In humans, the *CTNND1* gene is located at 11q11 and consists of 21 exons, of which exons 11, 18 and 20 are alternatively spliced. Inclusion of exon 11, which is predominantly neural, disrupts a nuclear localization signal (NLS), while exon 20 contains a nuclear export signal (NES)^26^. In addition, there are four additional isoforms of the protein, which vary in their transcriptional start sites. Of the four major isoforms, isoform 1 is abundant in mesenchymal cells, while isoform 3 appears preferentially expressed in epithelial cells^27–30^. The other two isoforms are less well characterized.

The p120 superfamily includes p120-catenin itself, δ-catenin (CTNND2) and ARVCF (<u>a</u>rmadillo repeat gene deleted in <u>v</u>elo<u>c</u>ardiofacial syndrome) all of which can compete for E-cadherin binding. Although it is unclear whether they substitute for one another in other cellular functions^31, 32^, evidence from animal studies suggests some compensatory roles. For instance, δ-catenin (CTNND2) knockdown phenotypes can be rescued with p120-catenin, and the combined depletion of δ-catenin and p120 generates more pronounced effects. However, levels of p120 are not altered by reducing δ-catenin protein levels^33^. In humans, *CTNND2* variants have been associated with autism spectrum disorders and other neurodevelopmental conditions^34–39^. Interestingly, the other p120 family member, *ARVCF,* lies in 22q11. While loss of *TBX1* in 22q11 is thought to cause the key malformations associated with velocardiofacial (VCF) syndrome [MIM: 192430], evidence from animal models suggests that *ARVCF* may also play a role in craniofacial development^40–43^.

Although both p120-catenin and its binding partner E-cadherin have been proposed as causative genes in non-syndromic palatal clefting and BCD syndrome^1–3^, the patients that we describe here present with a multisystem condition broader than the previously described p120-associated BCD cases. While our patients consistently possess palatal phenotypes (cleft or high arched palate) as well as eyelid and tooth anomalies, they also display additional features including severe hypodontia, cardiac, limb and neurodevelopmental anomalies. We hypothesize that these novel truncating variants in *CTNND1* affect both E-cadherin-dependent and -independent functions of p120-catenin, and, given the range of phenotypes seen in our cohort, should be considered more broadly to cause a VCF-like syndrome.

## Subjects and Methods

### Recruitment, consent and sample collection

Participants were recruited from one of following: South Thames Cleft Unit at Guy’s and St Thomas Trust (GSTT), London, UK; the University of Calgary, Alberta Children’s Hospital, Canada; from the Children’s Hospital of Philadelphia, USA; or, from the Deciphering Developmental Disorders (DDD) Study, United Kingdom (www.ddduk.org). *CTNND1* data access was specifically collected under DDD Project CAP180, focusing on cranial neural crest anomalies (ABM/KJL). All individual study protocols were approved by local Institutional Review Boards, including UK Ethics: GSTT (REC16/NI/0026, Northern Ireland REC), DDD (10/H0305/83, Cambridge South REC, and GEN/284/12, Republic of Ireland REC).

Medical and dental histories were taken, as well as detailed phenotyping by clinical geneticists with expertise in dysmorphology. Saliva for DNA extraction was collected from family trios using the Oragene® DNA (OG-500) kit. All patients also underwent high-resolution analysis for copy number abnormalities using array-based comparative genomic hybridization. Informed consent from all participants was obtained for publication of data and photographs in the medical literature. All families were offered genetic counseling.

### Exome sequencing and variant screening

Exome sequencing from trios was performed to identify gene variants. For patients recruited from DDD^44^, genomic DNA samples from trios were analysed at the Wellcome Trust Sanger Institute. Exome sequencing was performed using a custom Agilent SureSelect Exome bait design (Agilent Human All Exon V3 Plus with custom ELID # C0338371), 8-plex sample multiplexing and an Illumina HiSeq with 4 samples per lane and a mean depth of 50X. The exome analysis targeted 58.62 Mb of which 51.64 Mb consisted of exonic targets (39 Mb) and their flanking regions and 6.9 Mb consisted of regulatory regions. Alignment was performed using BWA1. Putative *de novo* variants were identified from trio BAM files using DeNovoGear5. Variants were annotated with the most severe consequence predicted by Ensembl Variant Effect Predictor (VEP version 2.6), and minor allele frequencies from a combination of the 1000 Genomes project (www.1000genomes.org), UK10K (www.uk10k.org), the NHLBI Exome Sequencing Project (esp.gs.washington.edu), Scottish Family Health Study (www.generationscotland.org), UK Blood Service and unaffected DDD parents. All flagged variants were automatically annotated with pathogenicity scores from two variant prioritisation algorithms (SIFT23 and PolyPhen24) and compared against the public Human Gene Mutation Database (HGMD) and the Leiden Open Variation Database (LOVD). For selected probands, Exome sequencing performed at the Yale Center for Genomic Analysis used genomic DNA isolated from saliva from the probands and their parents. The exons and their flanking regions of the genome were captured using IDT xGen exome capture kit followed by Illumina DNA sequencing (HiSeq 4000). Paired end sequence reads were converted to FASTQ format and were aligned to the reference human genome (hg19). GATK best practices were applied to identify genetic variants, and variants were annotated by ANNOVAR. Probands and parents were sequenced to a mean depth of 93-123 independent reads per targeted base across all the samples. In an average of 94.0% of targeted bases in all of the samples, the coverage was greater than 20X independent reads. Trio exome sequencing analysis on variants with allele frequency of less than 1% was carried out to identify *de novo* variants that are absent from the parents. Putative disease-causing variants were validated using whole genome amplified DNA, PCR and capillary sequencing.

### Mouse and Xenopus husbandry

Animal work was performed in accordance with UK Home Office Project License P8D5E2773 at King’s College London (KJL), University of Texas Southwestern Medical Center Institutional Animal Care and Use Committee protocols (DKM), the European *Xenopus* Resource Centre, Portsmouth UK, or the Yale University Institutional Animal Care and Use Committee protocols (MKK). Mice were genotyped according to standard procedures. Gestational ages for mice were determined by the observation of vaginal plugs, which was considered embryonic day 0.5 (E0.5) and further staging of animals according to Kaufman^45^. The following mouse strains were used: *Ctnnd1^fl/fl^* (MGI ID: 3640772)^8^*; β-actin::cre* (JAX strain 019099)^46^; and *Wnt1::cre* (JAX strain 022501)^47^. For each mouse experiment, a minimum of n=3 was examined unless otherwise noted. *X. tropicalis* embryos were produced by *in vitro* fertilization and raised to appropriate stages in 1/9MR + gentamycin as per standard protocols^48^. For *Xenopus* experiments, experimental numbers are stated in figures, with a minimum of n=30 in all experimental conditions.

### Human specimens

Human embryonic and fetal material was provided by the Joint MRC/Wellcome Trust (grant # 099175/Z/12/Z) Human Developmental Biology Resource (HDBR, http://www.hdbr.org) as whole embryos (Carnegie stage 13 (C13, day 28-32)) or sectioned embryos (Carnegie stage 21 (C21, day 50-52)).

### Generation of CTNND1 probe and mRNA in situ hybridization

A human *CTNND1* clone was identified from the Human ORFeome Collaboration^49^ (clone HsCD00513511), encoding *CTNND1* isoform 4, including the entirety of the armadillo repeats and the C-terminal domain. Probes made from this clone should recognize all four *CTNND1* transcripts. Digoxigenin-labeled antisense mRNA probes were produced by linearizing human *CTNND1* clones using BamH1 restriction enzyme, which produces a probe size of ∼900 base pairs, and *in vitro* transcription with the T7 High Yield RNA Synthesis Kit (E2040S) from New England Biolabs. *In situ* hybridization of mRNA on whole mount and paraffin embedded tissue sections was carried out as per standard protocols^50^, using an anti-digoxigenin-alkaline phosphatase coupled antibody.

### Immunofluorescent antibodies and staining

For immunostaining, mouse embryos at the indicated stages were fixed and processed according to standard protocols. Antigen retrieval was carried out in Tris-EDTA (pH 9) in a 90°C water-bath for 30 minutes. Primary antibodies used were: phospho-tyrosine p120-catenin clone 2B12, mouse mAb (1:150, Biolegend, Cat. No. 828301); delta 1 Catenin/CAS (phospho S-268) antibody [EPR2380], rabbit mAB (1:150, Abcam, Cat. No. ab79545); E-Cadherin [M168], mouse mAB (1:150, Abcam, Cat. No. ab76055); anti-E-cadherin (24E10), rabbit mAb (1:250, Cell Signaling Technology, Cat. No. 3195); rabbit anti-Pax2 Antibody (1:100, ThermoFisher Scientific, Cat. No. 71-6000); and mouse anti-Collagen Type II, clone 6B3 (1:50, MERCK, Cat. No. MAB8887). Secondary antibodies used were: Alexa Fluor® 488 (Invitrogen, A-11008), Alexa Fluor® 488 (Invitrogen, A-21204), Alexa Fluor® 546 (Invitrogen, A-11060), Alexa Fluor® 568 (Invitrogen, A-11011), Alexa Fluor® 594 (Invitrogen, A-21207), Alexa Fluor® 647 (Invitrogen, A-21235). All were diluted to 1:400 in phosphate-buffered saline (PBS) containing 0.5% Triton® X-100 (Sigma-Aldrich) and 1% bovine serum albumin. Slides were mounted in Fluoroshield Mounting Medium with DAPI (Abcam, ab104139) and cover slipped. *Xenopus* whole mount embryos and tadpoles were incubated with Hoechst (1:5000 of 20mg/ml, diluted in PBST). For hematoxylin and eosin (H&E) staining, slides were fixed, sectioned and stained according to standard protocols. Slides were then cover slipped with Neo-Mount (VWR, Cat. No. 1.09016.0500).

### Image acquisition

Images for sectional *in situ* hybridization experiments and for H&E slides were captured using a brightfield microscope (Nikon ECLIPSE Ci-L), with an attached camera (Nikon digital sight DS-Fi1) or with a NanoZoomer 2.ORS Digital Slide Scanner (Hamamatsu); NDP.view2 Viewing Software (U12388-01) was used to analyze the scanned images. Whole mount images of mouse pups and embryos, *Xenopus* and human embryos were captured using a Nikon SMZ1500 stereomicroscope with a Nikon digital sight DS-Fi1 (112031) camera. Fluorescent images of mouse palates and *Xenopus* epithelial cells were either acquired on a Leica SP5 confocal or Nikon A1R point scanning confocal; z-stacks of whole mount *Xenopus* tadpoles were captured by mounting the tadpoles on a Cellview Cell Glass Bottom Culture Dish (PS, 35/10 mm, CELLview™, Cat. No. 627860) in PBS. Image sequences were processed using the FIJI (Image J) analysis software.

### Micro-computed tomography (µCT)

For soft tissue scanning, mouse embryos were stained with a near isotonic 1% I2, 2% potassium iodine solution for 3 days and scanned to produce 6um voxel size volumes, using X-ray settings of 90kVp, 66uA and a 0.5 mm aluminium filter to attenuate harder X-rays. Camera binning was used to improve signal to noise ratios. For hard tissue staining, perinatal mice were scanned to produce 7.4um voxel size volumes using X-ray settings of 70kVp, 114uA and a 0.5 mm aluminium filter to attenuate harder X-rays. The specimens were analysed using Parallax Microview software package (Parallax Innovations Inc., Ilderton, ON Canada). Specimens were scanned using a Scanco µCT50 microcomputed tomographic scanner (Scanco, Brüttisellen, Switzerland). The specimens were immobilised in appropriately sized scanning tubes using cotton gauze.

### CRISPR/Cas9 knockouts in Xenopus tropicalis

The following non-overlapping single guide RNAs (sgRNAs) were designed to target *Xenopus tropicalis ctnnd1*: sgRNA1-CTAGCtaatacgactcactataGGAACGGGTGTGGGAGCCATgttttagagctagaa; sgRNA2 - CTAGCtaatacgactcactataGGGGTGGTATCCCACGCAAGgttttagagctagaa. sgRNA1 targets exon 3 and is thus predicted to disrupt isoform 1 only, while sgRNA2 targets exon 7 and is thus predicted to disrupt all four isoforms. Embryos were injected at the one or two cell stage and raised until indicated stages. For CRISPR/Cas9 experiments, statistical significance was defined as P<0.05 and analysed by chi-squared test or Fisher’s exact test.

## Results

### Identification of CTNND1 variants

Here, we identify 13 individuals with protein-truncating variants in *CTNND1.* Previously, all patients had undergone an array-based comparative genome hybridization analysis with normal results. A subset of patients had also been referred for other diagnostic tests, including 22q11 deletion, Down syndrome, CHARGE syndrome (*CHD7* sequencing), Noonan syndrome (*PTPN11* sequencing) and other conditions, but with no definitive diagnoses. Exome sequencing of the patients revealed eight novel variants in *CTNND1*, including six confirmed *de novo* variants (in 7 patients). Two individuals inherited their variant from affected parents while two other participants inherited a variant from a parent with a mild phenotype (Figure 1A). These truncating mutations included nonsense, splicing and frameshift variants (Table 1).

**Figure 1.**
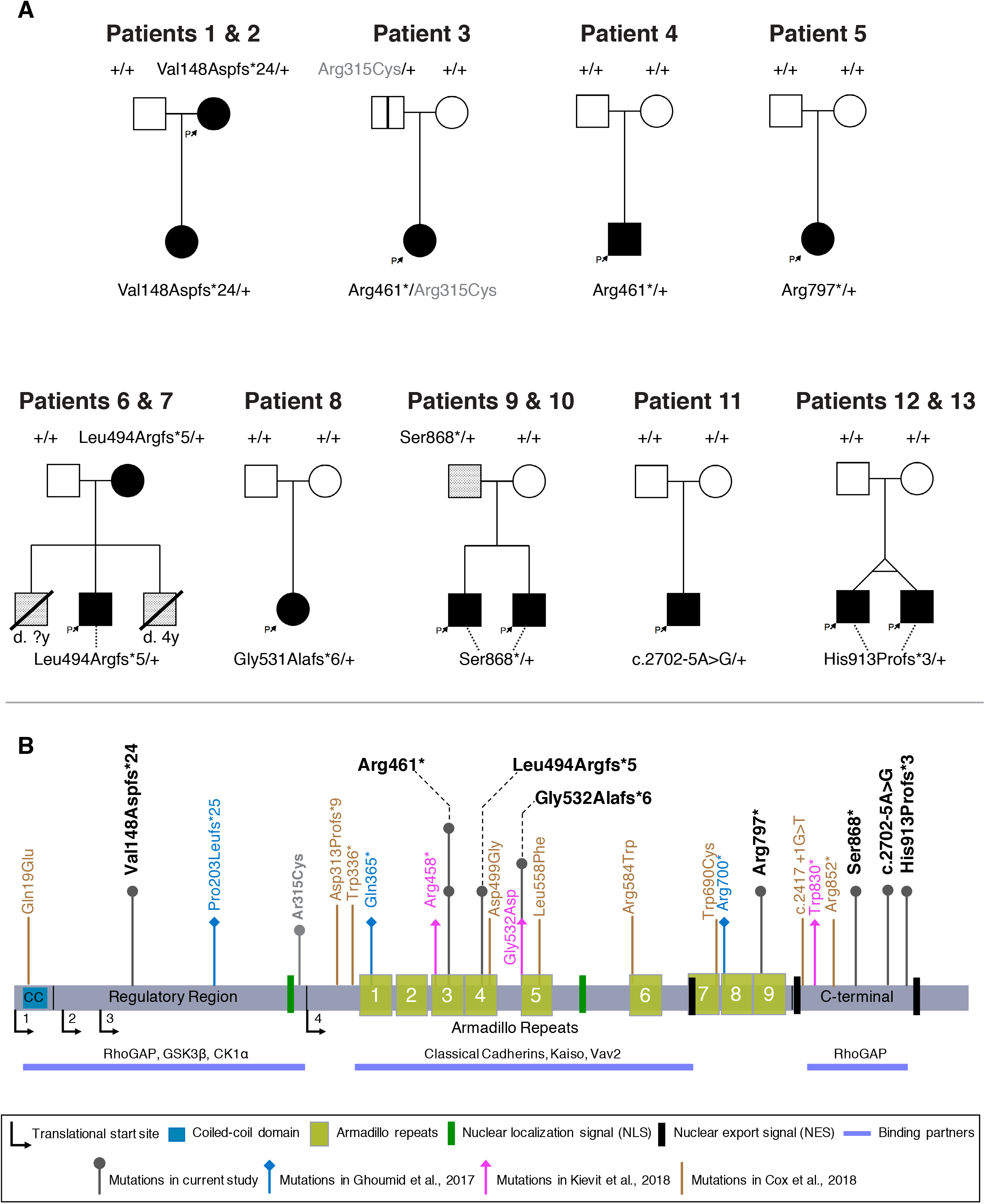
Pedigrees and identification of CTNND1 variants. [A] Pedigrees of individuals with identified variants. Filled boxes indicate affected individuals demonstrating collective phenotypes described in our cohort. A blank box with a vertical black line indicates an asymptomatic carrier (clinically unaffected). A box with an oblique line indicates a deceased individual. Lightly shaded boxes indicate individuals affected with one or more of the conditions described. [B] Schematic representation of the human p120-catenin protein structure and its domains. The variants described in our cohort are shown above the protein with a dark gray arrow. The light gray arrow with the (p.Arg315Cys) variant indicates the other *CTNND1* mutation found in Patient 3 which was inherited from the unaffected father [A]. Arrows in blue, pink and brown represent the variants and their locations reported in Ghoumid et al.^2^, Kievit et al.^1^ and Cox et al.^3^, respectively.

**Table 1:**
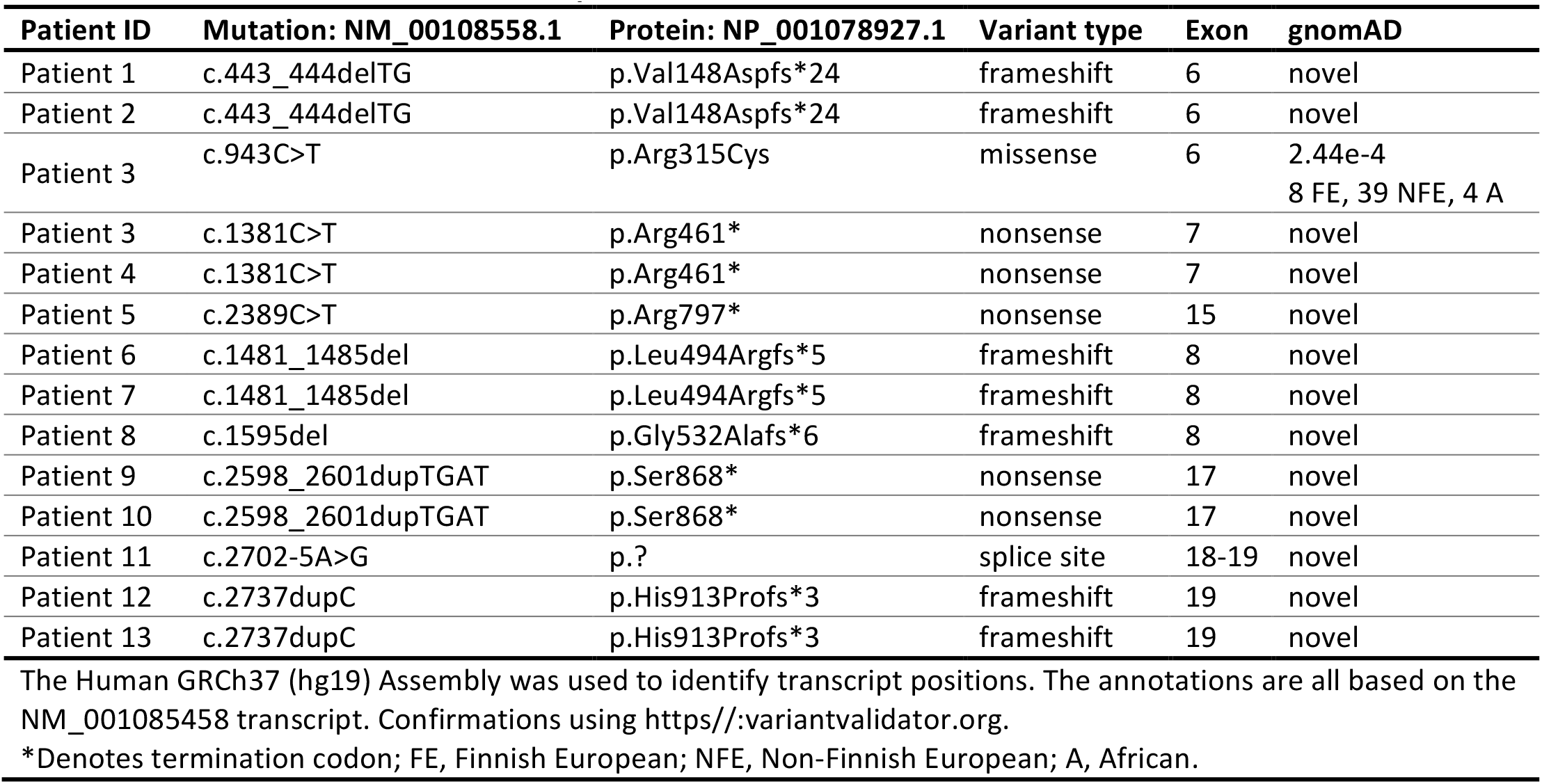
*CTNND1* variants in index patients.

| Patient ID | Mutation: NM_00108558.1 | Protein: NP_001078927.1 | Variant type | Exon | gnomAD |
| --- | --- | --- | --- | --- | --- |
| Patient 1 | c.443_444delTG | p.Val148Aspfs*24 | frameshift | 6 | novel |
| Patient 2 | c.443_444delTG | p.Val148Aspfs*24 | frameshift | 6 | novel |
| Patient 3 | c.943C>T | p.Arg315Cys | missense | 6 | 2.44e-4<br>8 FE, 39 NFE, 4 A |
| Patient 3 | c.1381C>T | p.Arg461* | nonsense | 7 | novel |
| Patient 4 | c.1381C>T | p.Arg461* | nonsense | 7 | novel |
| Patient 5 | c.2389C>T | p.Arg797* | nonsense | 15 | novel |
| Patient 6 | c.1481_1485del | p.Leu494Argfs*5 | frameshift | 8 | novel |
| Patient 7 | c.1481_1485del | p.Leu494Argfs*5 | frameshift | 8 | novel |
| Patient 8 | c.1595del | p.Gly532Alafs*6 | frameshift | 8 | novel |
| Patient 9 | c.2598_2601dupTGAT | p.Ser868* | nonsense | 17 | novel |
| Patient 10 | c.2598_2601dupTGAT | p.Ser868* | nonsense | 17 | novel |
| Patient 11 | c.2702-5A>G | p.? | splice site | 18-19 | novel |
| Patient 12 | c.2737dupC | p.His913Profs*3 | frameshift | 19 | novel |
| Patient 13 | c.2737dupC | p.His913Profs*3 | frameshift | 19 | novel |

The Human GRCh37 (hg19) Assembly was used to identify transcript positions. The annotations are all based on the NM\_001085458 transcript. Confirmations using https://variantvalidator.org.
\*Denotes termination codon; FE, Finnish European; NFE, Non-Finnish European; A, African.

*CTNND1* variants identified could be grouped according to the overall structure of the protein (Figure 1B). One variant falling within the N-terminal regulatory region was identified in Patient 1. Patient 1 has a *de novo CTNND1* c.443_444delTG (p.Val148Aspfs*24) mutation in exon 6. Targeted sequencing for this variant was carried out on the affected daughter (Patient 2), which segregates with the phenotypes in the family.

Four variants fell within the armadillo repeats, which are predicted to be crucial for interactions with E-cadherin. Two unrelated individuals (Patients 3 and 4) both had a *de novo* mutation in *CTNND1:* c.1381C>T (p.Arg461*) (Figure 1A-B). This variant results in a nonsense substitution and creates a stop codon in exon 7. In addition, Patient 3 had a rare variant in *CTNND1,* inherited paternally c.943C>T (p.Arg315Cys), which is present at a frequency of 2×10^-4^ in reference populations^51^. As the parent shares none of the phenotypes with the patient, this second variant is unlikely to be causative. Patient 5 had a *CTNND1* variant c.2389C>T (p.Arg797*) on exon 15. A *CTNND1* frameshift variant c.1481_1485del (p.Leu494Argfs*5) in exon 8 was identified in a mother and child; both are affected (Patients 6 and 7, respectively). In the same exon, Patient 8 had a *CTNND1* variant c.1594del (p.Gly532Alafs*6).

We found three variants affecting the C-terminal domain, present in five patients in three families. The variant c.2598_2601dupTGAT (p.Ser868*) was paternally inherited in a family with two affected siblings (Patients 9 and 10). The father is fit and healthy; however, his palate is narrow and high, and his nose is prominent. Patient 11 has a *de novo CTNND1* variant at the splice acceptor site of exon 19 designated as c.2702-5A>G, which is predicted to create a cryptic splice site, leading to a premature termination codon at the start of exon 19. Finally, Patients 12 and 13 are monozygotic twins carrying a *de novo* frameshift variant in *CTNND1*: c.2737dupC (p.His913Profs*3).

### Clinical presentation of patients with CTNND1 variants

Clinical phenotypes are summarized in (Table 2), and further details can be found in (Table S1). Photographs from participants show a number of shared craniofacial and oral features (Figure 2 and Figure 3, respectively) as well as other affected structures (eyes, ears and limbs (Figure S1)). Additional features including heart anomalies and neurodevelopmental conditions are noted in (Table 2 and Table S1).

**Figure 2.** Clinical presentation of individuals with a CTNND1 mutation. Facial photos (frontal and profile) show craniofacial features of patients. Note the narrow up-slanting palpebral fissures in Patients 3,4, 7-13; the hooded eyelids in patients 3, 4, 8-13; telecanthus in Patients 3,4,9-13; the high arched eyebrows in patients 1, 2, 6-8, 11-13 and the thin lateral eyebrows in Patients 1,5-11. Patients 1 and 4 had missing eyelashes medially from the inner canthus; Patients 1,2, 5 and 7 have distichiasis (double row of lashes), and mild ectropion of the lower eyelids were seen in Patients 1,5 and 6. As evident, no patient shows signs of hair sparsity. Most patients had wide nasal bridges with broad nasal tips while Patients 1,2, 8 and 11 were also diagnosed with congenital choanal atresia. Patients 1,2,7-9, 11 and 12 showed thin upper lips and while mid-face hypoplasia was observed, Patients 1,3,8,11 and 13 also had mandibular prognathism. Scars from cleft lip operations are seen in Patients 7, 9-13. Patient 3 was born with a submucous cleft palate, a bifid uvula and velopharyngeal insufficiency.

**Figure 3.**
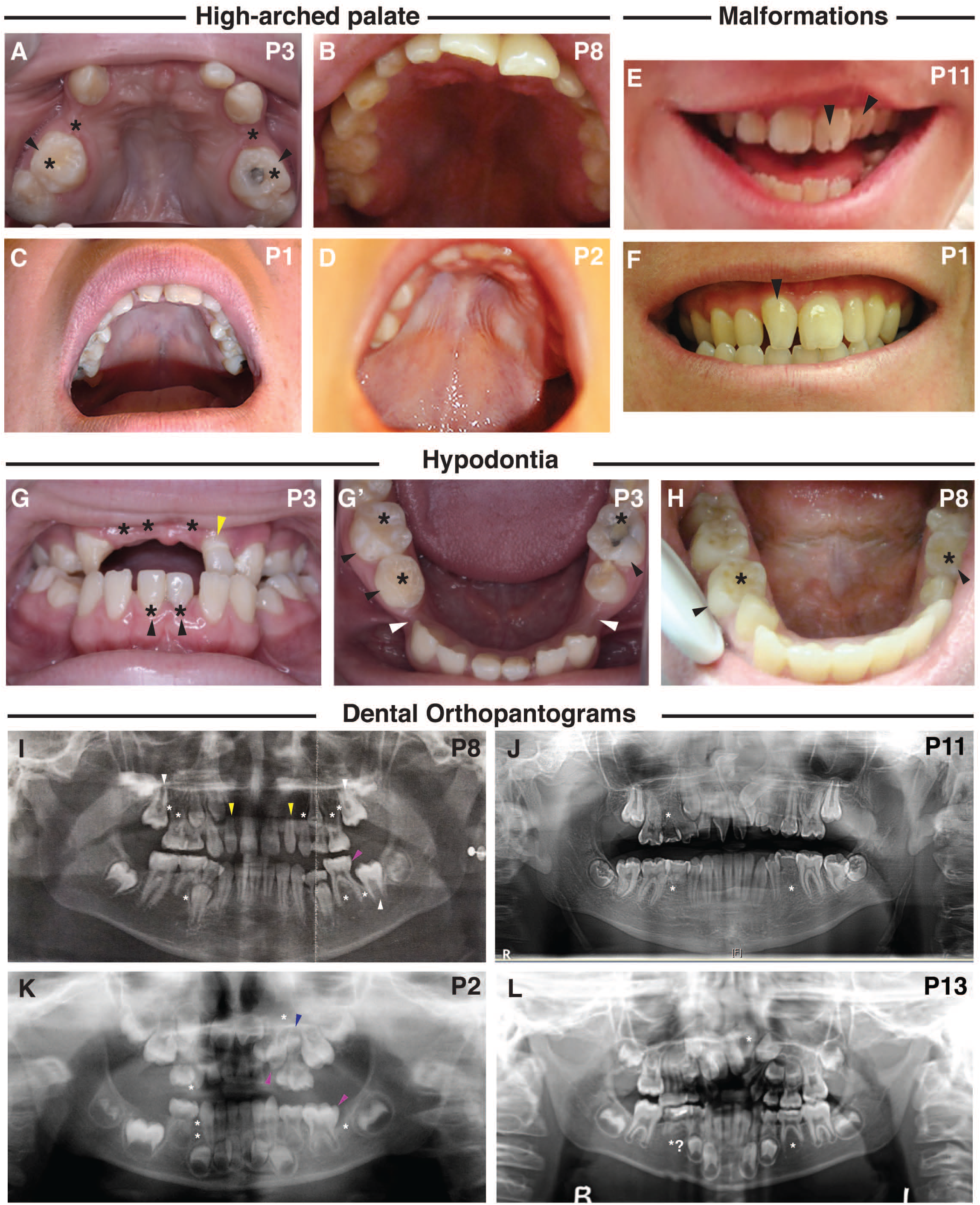
Dental manifestations and intra-oral phenotypes of patients with a CTNND1 mutation. [A-D] A high-arched palate was seen, shown are palates of Patients 1, 2, 3 and 8. [E-F] Abnormalities in the morphology of the dentition included: fissured incisors in Patient 11 [E, black arrowheads] and rotation of the incisors from the normal alignment shown in the non-cleft Patient 1 [F, black arrowhead]. [G-H] Hypodontia (tooth agenesis) was a common phenotype, indicated by the black asterisk. Black arrowheads indicate retained primary teeth. Patient 3 also has a diminutive upper left lateral incisor [G, yellow arrowhead] and wide inter-dental spacing [G’, white arrowheads]. [I-L] Dental orthopantograms (OPGs); missing teeth are indicated by white asterisks; diminutive teeth by yellow, macrodont teeth by magenta and supernumerary teeth by blue arrowheads, respectively. [I] OPG of Patient 8 at age 11, shows 8 missing permanent teeth (white asterisks) and shows the eruption of the second permanent molars (white arrowheads) in place of the missing first permanent molars. Also shown are diminutive upper right and left lateral incisors (peg-shaped) (yellow arrowheads), and a macrodont lower left second primary molar (magenta arrowhead). [J] OPG of Patient 11, at the age of 14, shows 3 missing permanent teeth (white asterisks), an ectopic maxillary left permanent canine and rotated maxillary centrals and left lateral incisors and dilacerated roots of the lower second permanent molars. [K] OPG of Patient 2, taken at 4 years, shows missing teeth including a missing lower left first permanent molar (white asterisks); a reported macrodont upper left primary canine (magenta arrowhead) with an underlying missing successor (white asterisk); a macrodont lower left second primary molar (magenta arrowhead) and a supernumerary tooth (blue arrowhead). [L] OPG for Patient 13, taken at 7.5 years, confirms absence of the upper left permanent lateral incisor and possibly the lower second permanent premolars.

**Table 2:**
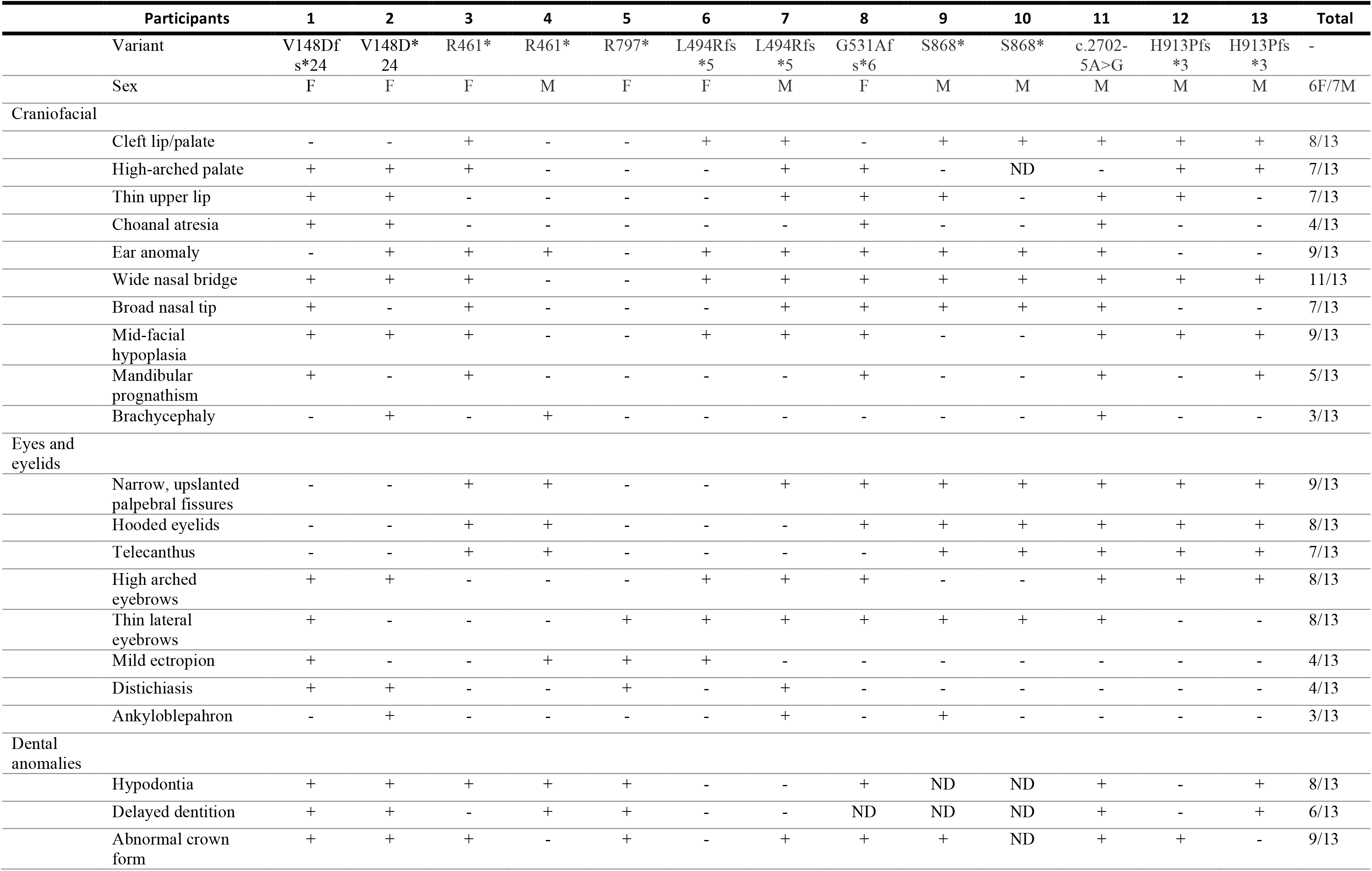
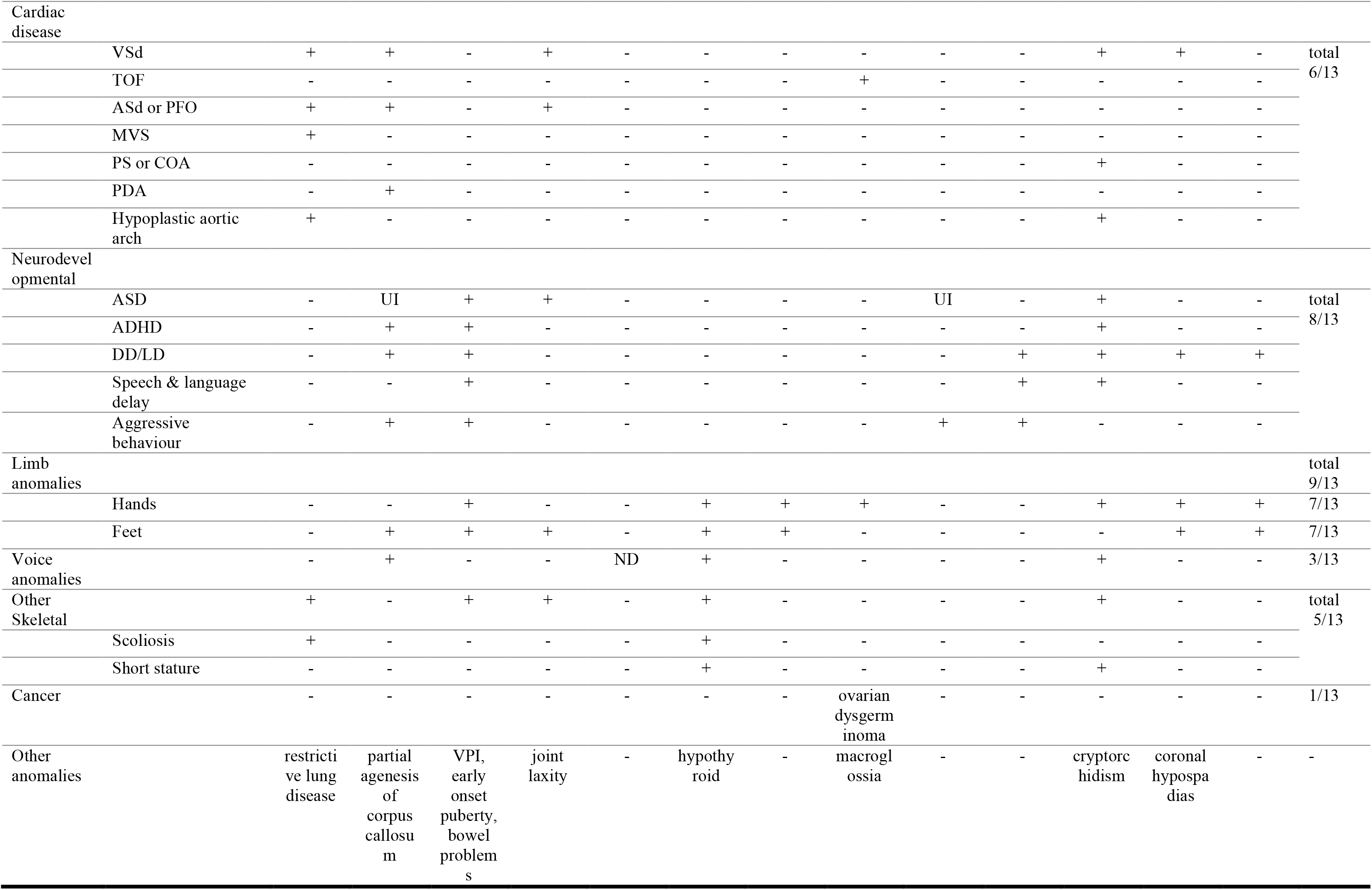
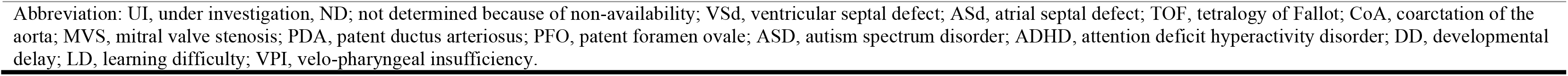
Clinical Summary of Individuals with *CTNND1 variant*.

Participants shared several distinctive eye features including short, up-slanted palpebral fissures (9/13), hooded eyelids (8/13), telecanthus (7/13), highly arched (8/13) and thin lateral eyebrows (8/13) and other eyelid anomalies such as nasolacrimal obstructions (1/13). These eye anomalies were clear from a young age (Figure S1A). A subset had ectropion (drooping lower eyelids, 4/13) and distichiasis (double eyelashes, 4/13). Many individuals had wide nasal bridges (11/13) with broad nasal tips (7/13), choanal atresia (4/13), either unilateral or bilateral atresia; malar flattening (mid-face hypoplasia) (9/13); mandibular prognathism (5/13); thin upper lips (7/13) and auricular abnormalities (9/13), particularly low-set ears and overfolded helices (Figure S1B).

Phenotypes with high penetrance involved oropharyngeal abnormalities including cleft lip and/or palate (CLP) (8/13), high-arched palate (7/13) or a combination of cleft and high-arched palate (Figures 3A-3D). A range of cleft sub-types was seen (Table S1). In addition, one participant had velopharyngeal insufficiency (VPI) and a bifid uvula. Of interest, three individuals presented with vocalization defects causing stridor and hoarseness or nasal speech.

Upon dental examination, all subjects were found to have intra-oral anomalies (Figure 3). In particular, congenital tooth agenesis (hypodontia) was frequently seen, with eight subjects missing between three and twelve adult teeth (Figure 3G-L; Table S2). Other anomalies included retained primary teeth and delayed eruption of the permanent teeth (6/13) (Table S1). Morphologic tooth anomalies were present, including diminutive permanent teeth/peg-shaped lateral incisors and fissured crowns of the permanent central and lateral incisors (Figures 3E-F; Table S1).

Beyond the craniofacial structures, the majority of the participants had limb and heart anomalies. Mild limb phenotypes (9/13) were present, including shorter fifth fingers, single transverse palmar crease, mild syndactyly between the 2,3 toes, sandal gaps and camptodactyly of the toes (Figure S1C). Congenital cardiac defects, which have not previously been associated with *CTNND1* variants, consistently occurred in our cohort. Six subjects had cardiovascular anomalies including tetralogy of Fallot, hypoplastic aortic arch, coarctation of the aorta, ventricular septal defect, atrial septal defect, mitral valve stenosis, patent ductus arteriosus and patent foramen ovale (Table 2 and Table S1). Finally, in addition to the craniofacial and cardiac anomalies, individuals presented with other phenotypes that added to the complexity of their conditions. Developmental delay and other neurodevelopmental problems were also observed (8/13). These often appeared from early toddler and school years and included mild learning difficulties, autism spectrum disorder, speech and language delay, and behavioral problems (Table S1). One individual was diagnosed with ovarian dysgerminoma stage III in the left ovary at the age of 12 years, which was treated with left oophorectomy followed by chemotherapy. Other infrequent anomalies included urogenital problems, scoliosis and partial agenesis of the corpus callosum (Table S1).

### P120 is expressed during human embryonic development

Although *P120* mRNA expression patterns have recently been documented during human and mouse palate development^3^, less is known about expression in the pharyngeal arch stages, which are likely to be important given the range of patient phenotypes. Therefore, we carried out mRNA *in situ* hybridization on human embryos using a probe that binds to all four C*TNND1* mRNA transcripts.

At Carnegie stage 13 (CS13), we found expression at multiple sites within the developing head, including the frontonasal processes, the forebrain, midbrain and rhombomeres (Figure 4B-4C). Robust expression was also detected in the maxillary and mandibular processes of the first pharyngeal arch (PA1), the second and third pharyngeal arches (PA2 and PA3, respectively) as well as in the proximal domains of the upper and lower limb buds (Figure 4A-4B). Signal was also weakly detected in the somites; however, strong expression was seen in the developing heart, trigeminal ganglion and the 10^th^ cranial nerve (Figure 4A-4B).

**Figure 4.**
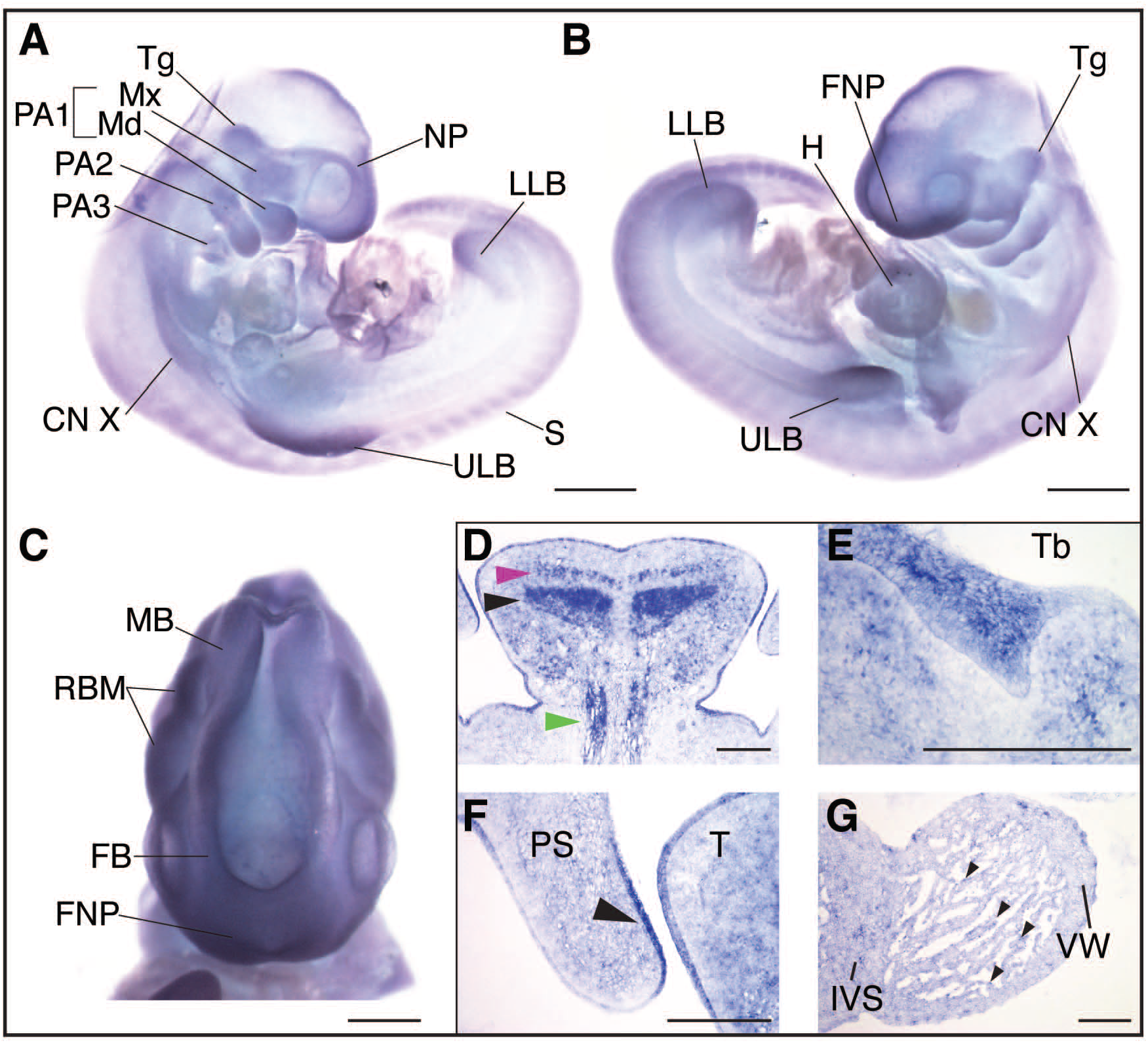
P120-catenin is expressed during relevant stages of human embryonic development. *CTNND1* mRNA *in situ* hybridization at human Carnegie stages 13 (CS13) [A-C] and 21 [D-G]. [A] Right lateral view of a CS13 human embryo, *CTNND1* mRNA is strongly expressed in the head in all three pharyngeal arches (PA1, PA2, PA3) and limb buds. Expression is specifically strong around the nasal placode and the maxillary and mandibular prominences. [B] Left lateral view, P120 is strongly expressed in the developing heart, frontonasal process, the trigeminal ganglion and the tenth cranial nerve. [C] P120 is ubiquitously expressed in the developing brain region in the rhombomeres, the forebrain and midbrain. [D-G] Coronal section through the head of a CS21 human embryo through a mid-palatal plane. [D] Strong expression is seen in the intrinsic muscles of the tongue: the superior longitudinal (magenta arrowhead), the transversal muscles of the tongue (black arrowhead) and the extrinsic genioglossus muscle (blue arrowhead). [E] *CTNND1* mRNA is strongly expressed in the epithelium of the developing tooth bud. [F] *CTNND1* is expressed on the dorsal epithelium of the palatal shelf (arrowhead) and in the epithelium of the tongue. [G] Expression is seen in the cardiomyocytes of the ventricular wall and the interventricular septum and in the cells of the endocardium (arrowhead). Scale bars = 100µm. Abbreviations: PA1, first pharyngeal arch; PA2, second pharyngeal arch; PA3, third pharyngeal arch; Tg, trigeminal ganglion; Mx, maxillary process; Md, mandibular process; CN X, tenth cranial nerve; ULB, upper limb bud; S, somites; LLB, lower limb bud; NP, nasal placode; H, heart, FNP, frontonasal process; Tb, mandibular tooth bud; PS, palatal shelf; T, tongue; IVS, interventricular septum; VW, ventricular wall.

By Carnegie stage 21, *CTNND1* mRNA was expressed in the brain (data not shown), tooth bud (Figure 4E), the epithelial lining of the tongue and oral cavity and in the tongue mesenchyme (Figure 4D). Expression was particularly strong in the intrinsic muscles of the tongue: the superior longitudinal and transversal muscles and in the extrinsic genioglossus muscle (Figure 4D). Moreover, expression was evident in the dorsal epithelial lining of the developing palatal shelves (Figure 4F). In the heart, *P120* expression was found in cardiomyocytes of the ventricular wall and interventricular septum, in addition to strong expression in the endocardium (Figure 4G). Expression was also found in the intrinsic epithelial lining of the stomach wall; both in the pyloric part of the stomach and in the inner walls of the stomach body, the pancreatic islets, the germinal center of the spleen, the epithelial lining of the bladder, hindgut and in the spinal cord and vertebral body (Figure S2).

### Expression of phosphorylated p120-catenin predicts fusion of the palatal seam

Because all of our participants had either cleft palate or associated palatal anomalies, we also assessed p120-catenin expression during palatal fusion in the mouse, which occurs from embryonic day 12.5 (E12.5) to E15.5 (Figure 5A-5D). To examine this, we used two antibodies recognizing phosphorylated forms of p120-catenin: a tyrosine-phosphorylated form, or phosphorylation at serine 268 (pS-268), which is proposed to trigger disruption of epithelial cadherin-catenin complexes^52, 53^. Neither of these forms of p120-catenin had been previously analyzed in the palate. In palatal cross-sections at E14.5, the medial epithelial seam (MES) is evident (Figure 5B), followed a few hours later with dissolution of the seam at E14.75 (Figure 5C). While E-cadherin is expressed as expected in the MES^54^ (Figure 5F, J), the two forms of p120-catenin show very distinctive distributions. As the seam undergoes EMT, at E14.5, pS-268 is strongly expressed as predicted in cell-cell interfaces of the periderm layer along the medial seam, clearly co-localising with E-cadherin (Figure 5E-5F). As the seam degrades, E-cadherin expression is lost while p120-catenin expression remains (Figure 5G-5H, white arrowheads). To our surprise, we find phospho-tyrosine p120 staining in both the mesenchymal and the epithelial cells, with a clear enrichment marking the border between the epithelial and mesenchymal populations (Figure 5I-5J, pink arrowheads). This distribution appears unique to this stage of palate formation consistent with reports that p120-catenin is tyrosine phosphorylate in an EGFR-dependent manner^55^, and continues during degradation of the seam while E-cadherin expression decreases (Figure 5K-5L, pink arrowheads). As a control, in earlier stages (E11-12.5), the phospho-tyrosine expression is much lower and nearly identical to the pS-268 staining (data not shown).

**Figure 5.**
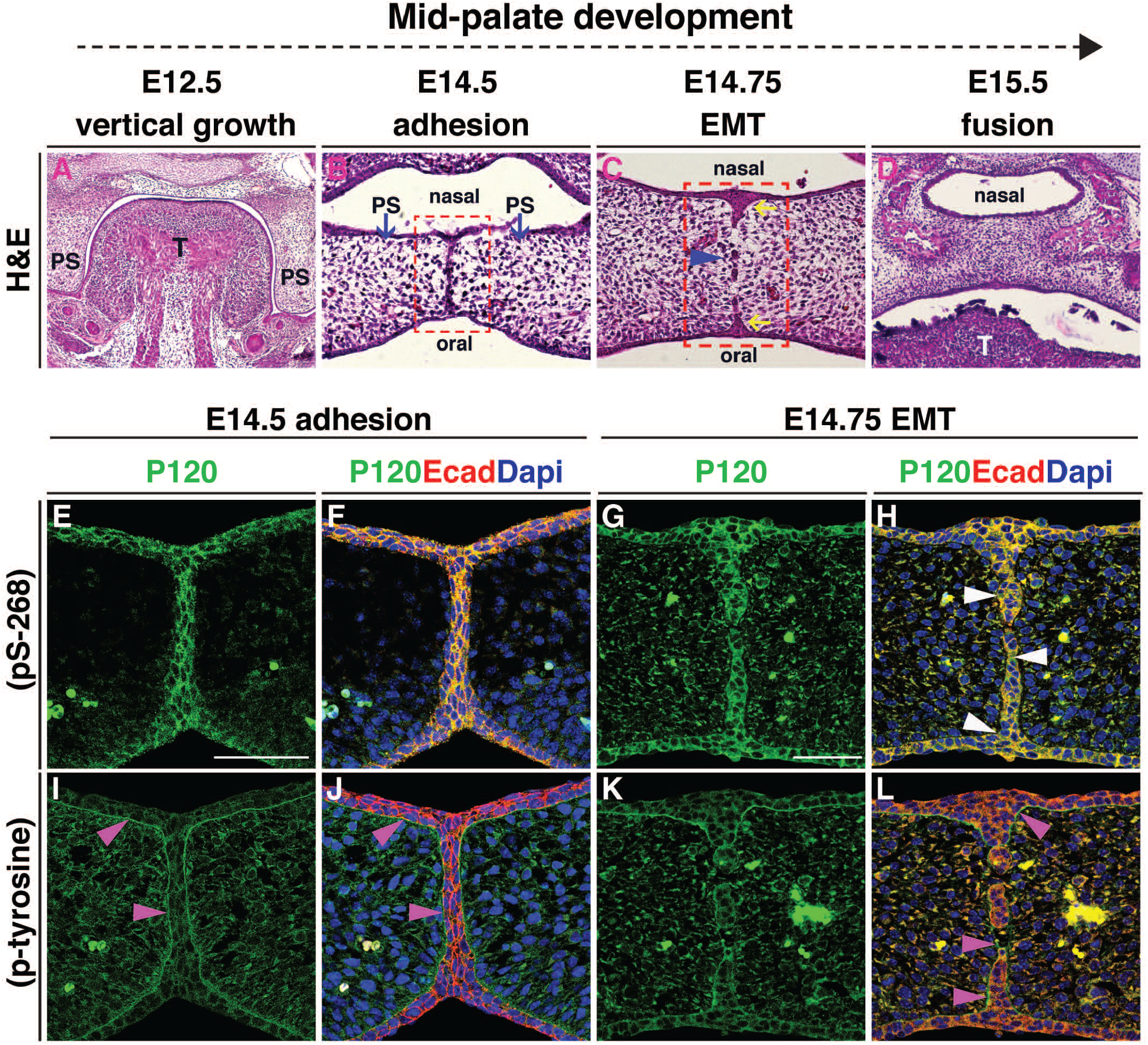
Expression of phosphorylated p120-catenin predicts fusion of the palatal seam. [A-L] All images are coronal sections of CD1 wild-type murine embryos at consecutive stages of palatal development. [A-D] Hematoxylin and eosin (H&E) staining illustrates successive stages of palatogenesis from embryonic day (E) 12.5 to E15.5. [B] At E14.5, following horizontal elevation, the opposing palatal shelves (blue arrows) meet and adhere to form the medial epithelial seam (MES). [C] EMT occurs at E14.75 when the MES breaks down, forming epithelial islands (blue arrowhead); the nasal and oral epithelial triangles form (yellow arrows). [D] At E15.5 palatal shelves are fused. Red box in [B] marks the regions shown in [E-F, I-J]. Red box in [C] marks the regions shown in [G-H, K-L]. [E-L] Immunofluorescent staining for either pS-268 or p-tyrosine p120-catenin antibodies (green) shown independently in [E, G, I, K], or in a merge with E-cadherin antibody staining (red) and DNA/DAPI stain (blue) [F, H, J, L]. [E-F, I-J] At E14.5, both forms of p120-catenin are expressed, with pS-268 strongly expressed in the periderm at the midline seam co-localizing with E-cadherin [E-F], while p-tyrosine clearly enriched in the area marking the border between the epithelial and mesenchymal populations [I-J, pink arrowheads]. [G-H, K-L] At E14.75, pS-268 p120-catenin is strongly expressed in the epithelial islands and the oral and nasal epithelial triangles; this is co-localised with E-cadherin during EMT and endocytosis while p120-catenin expression remains in some areas [H, white arrowheads]. In contrast, p-tyrosine p120-catenin expression surrounds E-cadherin positive epithelial islands, while E-cadherin expression has disappeared in the intervening mesenchymal cells (L, pink arrowheads). Scale bars = 50µm. Abbreviations: T, tongue; PS, palatal shelf.

### Heterozygous loss of p120-catenin leads to structural changes in the laryngeal apparatus

Some of our participants presented with anomalies associated with dysfunction of their velopharyngeal muscles and voice irregularities (Table S1 and Table 2), a phenotype described in patients with velocardiofacial syndrome^56–58^. Antibody staining confirmed presence of p120-catenin protein during development of the laryngeal and pharyngeal tissues in the mouse (Figure S3A). We then examined the laryngeal structures of mutant mice compared to their littermate controls at E16.5, P1 and P2.5 (Figure 6). To do this, we crossed a mouse carrying the ubiquitous *β-actin::cre* driver with *Ctnnd1^fl/fl^* mice in order to generate heterozygous mutants^59, 60^ (Figure 6C, 6H, 6M, 6R). Because we previously showed that the vocal ligaments originated from the neural crest^61^, we also generated tissue-specific *Ctnnd1* heterozygotes using the neural crest specific driver, *Wnt1::cre*^62^ (Figure 6E, 6J, 6O). We found identical laryngeal anomalies in the heterozygous mutants in both mouse crosses, confirming the neural crest-specificity of these phenotypes.

**Figure 6.**
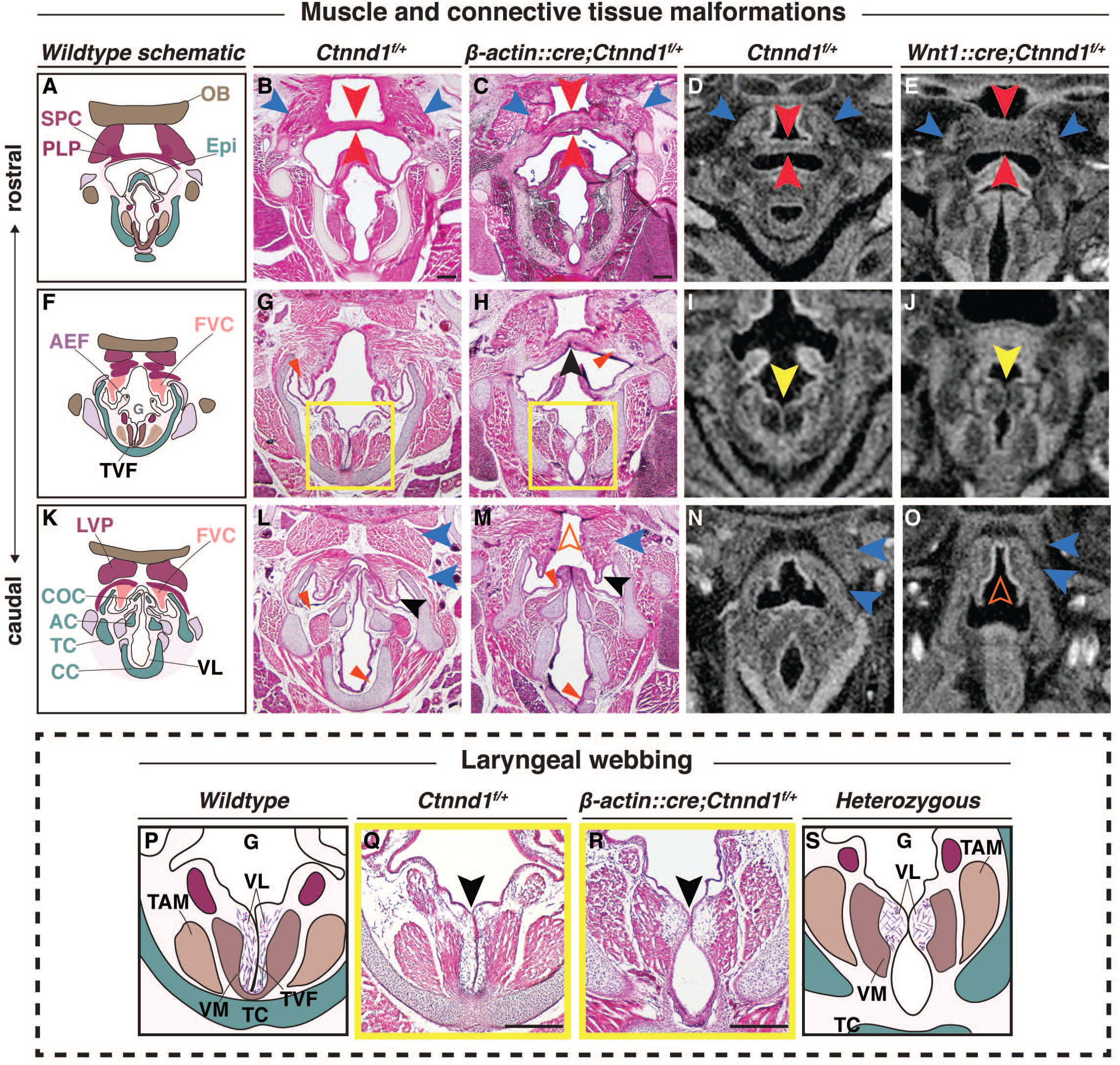
Heterozygous loss of p120-catenin leads to structural changes in the laryngeal apparatus. [A-O] Progression of the pharyngeal and laryngeal anomalies. [A, F, K] Schematics show the organization of the wildtype oropharynx from the more rostral (A) to caudal (K) planes. Haematoxylin and eosin (H&E) staining of coronal sections through control [B, G, L: *Ctnnd1^fl/+^*] and heterozygous mutants [C, H, M: *β-actin::cre/+; Ctnnd1^fl/+^*] littermate at postnatal stage (P1). [B-C] The SPC (blue arrowhead) and PLP (red arrowhead) in mutants are disorganized with an increased thickness in the PLP cranio-caudally [C] as compared to the controls [B]. [G-H] The FVC (vestibular folds) are well-defined in the controls with abundant ligaments [G, red arrowhead]. The FVC are fused in the mutant mice [H, black arrowhead] with ill-defined vestibular ligaments (H, red arrowhead). [L-M] The muscle attachments (blue arrowheads) superior to the FVC (black arrowhead) are well organized bilaterally in the controls surrounding the COC [L]. Caudally, when the FVC separated in the mutants it appeared hypoplastic (black arrowhead) as did the COC. The muscles (blue arrowheads) were ectopically fused to the LVP, producing an appearance of a ‘high-arched’ epiglottal area [M, orange hollow arrowhead]. ***[D-E, I-J, N-O] Neural crest specific mutants showed comparable laryngeal phenotype.*** Microcomputed tomographic (µCT) soft tissue scans of E16.5 control [D, I, N: *Ctnnd1^fl/+^*] or neural-crest specific [E, J, O: *Wnt1::cre/+; Ctnnd1^fl/+^*] heterozygous mutant littermates. [D-E] Compare the PLP in control [D] to the very thick PLP muscle seen in mutant [E, red arrowheads]. Compare the SPC in control [D] to the disorganized and hypoplastic SPC muscles seen in mutants [E, blue arrowheads]. [I-J] Laryngeal webbing was observed in mutant TVF [J, yellow arrowhead] compared to parallel TVF in control littermate [I, yellow arrowhead]. [N-O] Note aberrant muscle attachments (blue arrowheads) in [O] compared to control [N]. Control [N] epiglottal region compared to the high-arched epiglottal area observed in mutant littermate [O, orange hollow arrowhead]. ***[P-S] The laryngeal webbing phenotype.*** [P and S] Schematic representations of the wild-type [P] and mutant [S] anatomy at the vocal folds (TVF) from yellow-boxed insets in [G] and [H], respectively. [Q-R] H&E staining of coronal sections through control [Q: *Ctnnd1^fl/+^*] and heterozygous mutant [R: *β−actin::cre/+;Ctnnd1^fl/+^*] littermate at P1. [Q] In controls, well-defined vocal ligaments (VL) run parallel to the true vocal fold/cords (TVF). Underlying, the vocalis muscle (VM) and the thyroarytenoid muscle (TAM) are clearly attached and well-organised. [R] Laryngeal webbing is seen in the heterozygous mutant mice, where the vocal ligaments (VL) accumulate at a thin contact point (black arrowhead) thus perturbing the correct muscle attachments of the VM and TAM. Scale bars = 100µm. Abbreviations: SPC, Superior Pharyngeal Constrictor; PLP, Palatopharyngeus Muscle; TAM, Thyroarytenoid Muscle; VM, Vocalis Muscle; LGF; HB, Hyoid Bone; Epi, Epiglottis; OB, Occipital Bone; LVP, Levator Veli Palatini Muscle; AEF, Aryepiglottic Fold; TVF, True Vocal Fold; VL, Vocal Ligament; FVC, False Vocal Cord; CC, Cricoid Cartilage; TC, Thyroid Cartilage; AC, Arytenoid Cartilage; COC, Corniculate Cartilage.

Specifically, in control *Ctnnd1^fl/+^* mice, the palatopharyngeus (PLP) muscle, which elevates the larynx, is well defined and runs uniformly perpendicular to the epiglottis thereby attaching to the superior pharyngeal constrictor muscle (SPC) on either side (Figure 6A, 6B and 6D). On the other hand, the PLP and the SPC were both severely disorganized in both sets of heterozygous mice with an apparent increase in the cranio-caudal thickness of the PLP muscle (Figure 6C and 6E). Second, a striking phenotype known as laryngeal webbing was observed (compare controls, Figure 6G, 6I, 6Q to mutants Figure 6H, 6J, 6R). Typically, the bilateral vocal cords are parallel and meet at the midline (Figure 6F-6G, with inset schematized and shown in 6P and 6Q). The outer layer of the vocal fold is made of an epithelium that encapsulates the lamina propria comprising the vocal ligaments (Figure 6P and 6Q). These two layers function as the vibratory components for phonation and oscillation. Instead, in heterozygous mutant mice, the vocal ligaments show only a brief contact point between the opposing epithelia (Figure 6H, with inset schematized and shown in 6R and 6S). The vocal cords are also thinner, lacking the lamina propria (Figure 6R). Laryngeal webbing was also seen in the *Wnt1::cre* heterozygotes (Figure 6J) compared to their littermate controls (Figure 6I).

While the vestibular folds were well demarcated and the ligaments within them clearly defined in controls (Figure 6G), the vestibular folds in the heterozygous mice were ectopically fused and the ligaments sparse and dispersed (Figure 6H). Caudally, where the vestibular folds surrounded the normal corniculate cartilage (COC) (Figure 6 K, L); the folds have separated in the *Ctnnd1* heterozygotes, albeit hypoplastic (Figure 6M). Similarly, the COC appeared hypoplastic and devoid of the underlying lamina propria (Figure 6M). Finally, in mutants, the muscles were ectopically fused to the levator veli palatini muscles, which were then fused to the cranial base (Figure 6M). This, in turn, gave the impression of a high-arched epiglottal area; a defect also found in the *Wnt1::cre* heterozygous mutants (Figure 6O).

We also explored other craniofacial phenotypes in our heterozygous mouse model. Compared to their littermate controls (Figure S3B, a-e), mutant mice did not show any cleft lip (Figure S3B, f), face or limb dysmorphologies (Figure S3B, f-h) or cleft palate (Figure S3B, i) (n=12). This was confirmed by micro-computed tomography (µCT) to check for associated bony defects (n=6) (Figure S3B, j).

### P120-catenin isoform 1 function is required in multiple organ systems

While genetic mutation of *p120-catenin* in mouse models revealed a role for the neural crest in oropharyngeal development, analysis of multi-system involvement of p120-catenin was difficult due to embryonic lethality of the homozygous null mice^5, 9^. We therefore turned to the frog *Xenopus,* where *in vivo* function of p120-catenin has been well studied^11, 12, 63^. Previous analyses of p120-catenin requirements were mainly performed with antisense morpholino oligonucleotide (MO) knockdowns, which transiently prevent protein translation^11^. Instead, to create genetic mutants, we used CRISPR*/*Cas9 approaches, allowing us to specifically delete different p120-catenin isoforms^64^. As noted in the introduction, isoform 1 (full length at 968 amino acids (aa)) is most abundant in mesenchymal cells, while isoform 3 (start at aa 102) is preferentially expressed in epithelial cells^27–30^. Isoforms 2 and 4, which start at 55 aa and 324 aa, respectively, are less well characterized.

Embryos were injected at the one cell stage with single guide RNAs (sgRNAs) targeting either of two coding exons, exon 3 or exon 7 (sgRNA1 and sgRNA2 respectively, Figure 7A). Disruptions in exon 3 are predicted to only affect isoform 1, while sgRNA2 targeting exon 7 disrupt all four isoforms.

When embryos were scored at gastrula stages following sgRNA1 injections, disrupted or delayed blastopore closure was evident (n=30/42 vs. 2/30 in the controls) (Figure 7B). Furthermore, we noted severe early lethality (Figure 7D), especially using sgRNA2 which blocked all isoforms (Figure 7D). Notably, by neurula stages the majority of these mutants died due to a loss of integrity in the epithelium (data not shown).

Since the most well-established epithelial role for p120-catenin is in complex with E-cadherin at cell-cell junctions, we first examined E-cadherin localization in the neurectoderm at stage 11, as gastrulation was concluding. Indeed, in uninjected controls, high levels of p120-catenin and E-cadherin were found co-localized at the cell interface (Figure 7C, a-d). E-cadherin is expressed throughout the cell membrane (Figure 7C, b), whereas p120-catenin, though localized to the cell membrane, appears distributed in puncta (Figure 7C, a). Upon p120-catenin deletion, the expression levels of endogenous E-cadherin in the epithelial cells was diminished particularly at the interface between the cells, leaving only spot-like localization of both proteins at the tricellular junctions of these epithelial cells (Figure 7C, e-h). The residual expression of p120-catenin may be due to maternal loading of the protein, as the CRISPRs should only affect zygotic transcription, or due to mosaicism of the CRISPR deletion.

As the sgRNA2 CRISPR was predicted to disrupt all four isoforms and led to severe lethality by neurula stages, the majority of analyses were performed using the sgRNA1 CRISPR, which is predicted to disrupt the predominantly mesenchymal isoform 1. A proportion of the knockout animals survived past the neurula stages, possibly due to mosaicism, and were examined at stage 46 to determine whether craniofacial and organ development had occurred normally. We observed obvious craniofacial defects in the CRISPR mutants, including a reduction in the width and height of the head (Figure 7E, l-n), a hypoplastic mouth opening (Figure 7E, m), delayed breakdown of the cement gland (Figure 7E, l-m), heart and gut looping anomalies (Figure 7E, n). Following on from the disorganization of the laryngeal muscles seen in the mouse mutants (Figure 6), antibody staining against Pax2 was used to label the muscle fibers while anti-collagen 2 (col2) antibody labelled craniofacial cartilages in the mutants (Figure 8A, a-h). In control animals, the muscle fibers were well-organised and straight while in the mutants, the muscle morphology appeared disorganized, particularly the rectus abdominus muscle, with muscle striations being replaced by irregularly shaped fibers (Figure 8A, f-g). Consistent with previous observations (Figure 7), craniofacial cartilages were hypomorphic, and compacted both in the anterior-posterior and dorsal-ventral axes (Figure 8A, a and e). However, morphology of the chondrocytes appeared normal (Figure 8A, d and h).

Finally, since the participants (6/13) had a high frequency of congenital heart defects and because p120 is strongly expressed in the heart of human, mouse and frog embryos, we examined the hearts in the CRISPR-knockout tadpoles. Notably, the strong expression of p120 seen in the different heart chambers in the control tadpoles was lost when p120 was knocked down (Figure 8B, p). The majority of mutant tadpoles had heart anomalies including heart-looping defects (Figure 7E, n; Figure 8B, n). Notably, E-cadherin is not expressed in the normal heart or the muscles (Figure 8B, I), suggesting that the heart and muscle phenotypes may be manifestations of E-cadherin independent functions of p120.

## Discussion

This work expands upon the spectrum of abnormalities associated with *CTNND1* variants beyond non-syndromic cleft lip/palate (CLP) and BCD^1–3^. Most notably, we describe in detail characteristic craniofacial features including choanal atresia and unusual patterns of hypodontia as well as heart, limb, laryngeal and neurodevelopmental anomalies. We find expression of *CTNND1* mRNA during development of the pharyngeal arches in human embryos and we define the profile of two phosphorylated forms of p120 in the mouse palate. Finally, genetic approaches in mouse and *Xenopus* demonstrated novel roles for *CTNND1* in the oropharynx, craniofacial cartilages and in the heart. Thus, our data implicate *CTNND1* variants as causative of a broad-spectrum syndrome that overlaps with DiGeorge velocardiofacial syndrome as well as other disorders of craniofacial development such as CHARGE and Burn McKeown syndromes^65–68^. All of these syndromes could be collectively considered to be neurocristopathies. Notably, the neural crest specific disruption of *CTNND1* in our animal models supports this role for *CTNND1* as a candidate neurocristopathy gene and we suggest that these newly identified variants likely highlight both epithelial and mesenchymal roles for p120-catenin.

Prior to our study, the majority of the participants did not have a recognizable or a diagnosed condition when they were recruited. Here, we demonstrate that they collectively share consistent characteristic phenotypic features that suggest that mutations in *CTNND1* may lead to a much broader phenotypic spectrum than previously described^1, 2^. For instance, low set ears were reported in one case of BCD by Kievit and colleagues^1^; we find multiple participants with auricular anomalies particularly the low-set ears and over-folded helices (Figure S1B, Table S1). Similarly, syndactyly was reported in one of the *CTNND1* patients described in Ghoumid et al.^2^, and clinodactyly (one patient) and camptodactyly (two patients) were reported by Kievit et al^1^. Again, we find limb anomalies consistently associated with *CTNND1* variation (Figure S1C, Table S1). The cardinal features of BCD include ectropion of the lower eyelids, euryblepharon and lagopthalmos^69, 70^; while five of our patients showed these eye manifestations (Figure 2; Table 2), we also found short up-slanting palpebral fissures, hooded eyelids, high arched eyebrows and telecanthus (Figure S1A, Table 2 and Table S1). As BCD is associated with both *CTNND1* and *CDH1* (E-cadherin) variants, some of these phenotypes may represent distinctive functions of the E-cadherin-p120 complex; the majority of these functions could be attributed to a role for the cadherin-catenin in epithelia^71^.

Of note, eight individuals had severe hypodontia, including missing permanent canines and first permanent molars, even in those without cleft lip/palate. Thus, missing canines and molars could be classified as a microform cleft anomaly, especially when found in association with high-arched palate^72^ (Figure 3, I and K; Table S2).

Beyond the known phenotypes associated with *CTNND1* and *CDH1*, we note the novel phenotypes seen in our patients, which include the heart anomalies and behavioral disorders. These have not been reported previously in patients with a BCD diagnosis. Nevertheless, our findings suggest that both *CTNND1* and *CDH1* should be tested in patients with congenital orofacial and cardiac anomalies. A key finding was choanal atresia in four individuals; given the rarity of this anomaly, both *CTNND1* and *CDH1* should be considered during genetic profiling of patients with this anomaly, in addition to CHARGE and other syndromes noted above. Indeed, Nishi et al. (2016) reported cleft lip, right choanal atresia, a congenital cardiac anomaly (tetralogy of Fallot), agenesis of the corpus callosum, upslanted palpebral fissures and ear anomalies in a patient with *CDH1* mutation^73^; however, at the time, this was not diagnosed as BCD.

While all of the variants found in the present study resulted in truncations of p120-catenin, they fell broadly into three distinct groups: those falling within the N-terminal regulatory region (p.Val148Aspfs*24), those disrupting the armadillo repeat region and presumably subsequent interactions with E-cadherin (e.g., p.Arg461*, p.Arg797*, p.Leu494Argfs*5 and p.GLy532Alafs*6), and those falling in the C-terminal domain (p.Ser868*, the splice variant c.2702-5A>G and p.His913Profs*3). Interestingly, those probands with C-terminal truncations had the most complete cleft lip and palate phenotypes. This was consistent with previous reports by Kievit et al.^1^ who reported a nonsense mutation (p.Trp830*) and Cox et al.^3^ who reported p.Arg852* and a splice site mutation (c.2417+G>T)^3^. As these C-terminal truncations would all be predicted to retain E-cadherin binding, but lose crucial RhoGAP interactions^24^, one might hypothesize that a mutation in this region prevents p120 clearing from the epithelial complex, which is necessary for seam dissolution during palate closure. Therefore, future analyses should focus on whether these C-terminal truncations are acting in a dominant-negative manner, and preventing clearance of E-cadherin from the seam.

With regards to non-epithelial functions of p120, some of the phenotypes that this study, and others, have reported, could be explained by the known interactions of p120 in the Wnt signaling pathway^20^. Epithelial-specific knockouts of p120 (using a *keratin-14* promoter) did not show tooth agenesis^10^, suggesting that the tooth anomalies in our patients do not arise from the epithelial functions of p120. In support of this, two key genes implicated in tooth agenesis are the Wnt ligand, *Wnt10A* and a Wnt target gene *Axin2*^74–78;78–84^. The Wnt signaling pathway may also explain the laryngeal findings (Figure 6), as knockout of the Wnt transducer β-catenin is also known to lead to similar vocal fold anomalies^85^ as those seen in our neural crest specific *p120-catenin* heterozygotes (Figure 6). Furthermore, knockout of the mesenchymal form of p120 (isoform 1) in *Xenopus* (Figure 7 and Figure 8), confirm prior studies on p120-catenin in the neural crest, where the p120-catenin association with Wnt signaling is well-established^32, 86, 87^. Thus, we hypothesize that a subset of p120 phenotypes can also be attributed to Wnt perturbation in the neural crest (Figure 9). The heart defects seen in our patients could also be attributed to a failure in neural crest development, which is known to be crucial for development of the septum and valves^88–92^.

**Figure 7.**
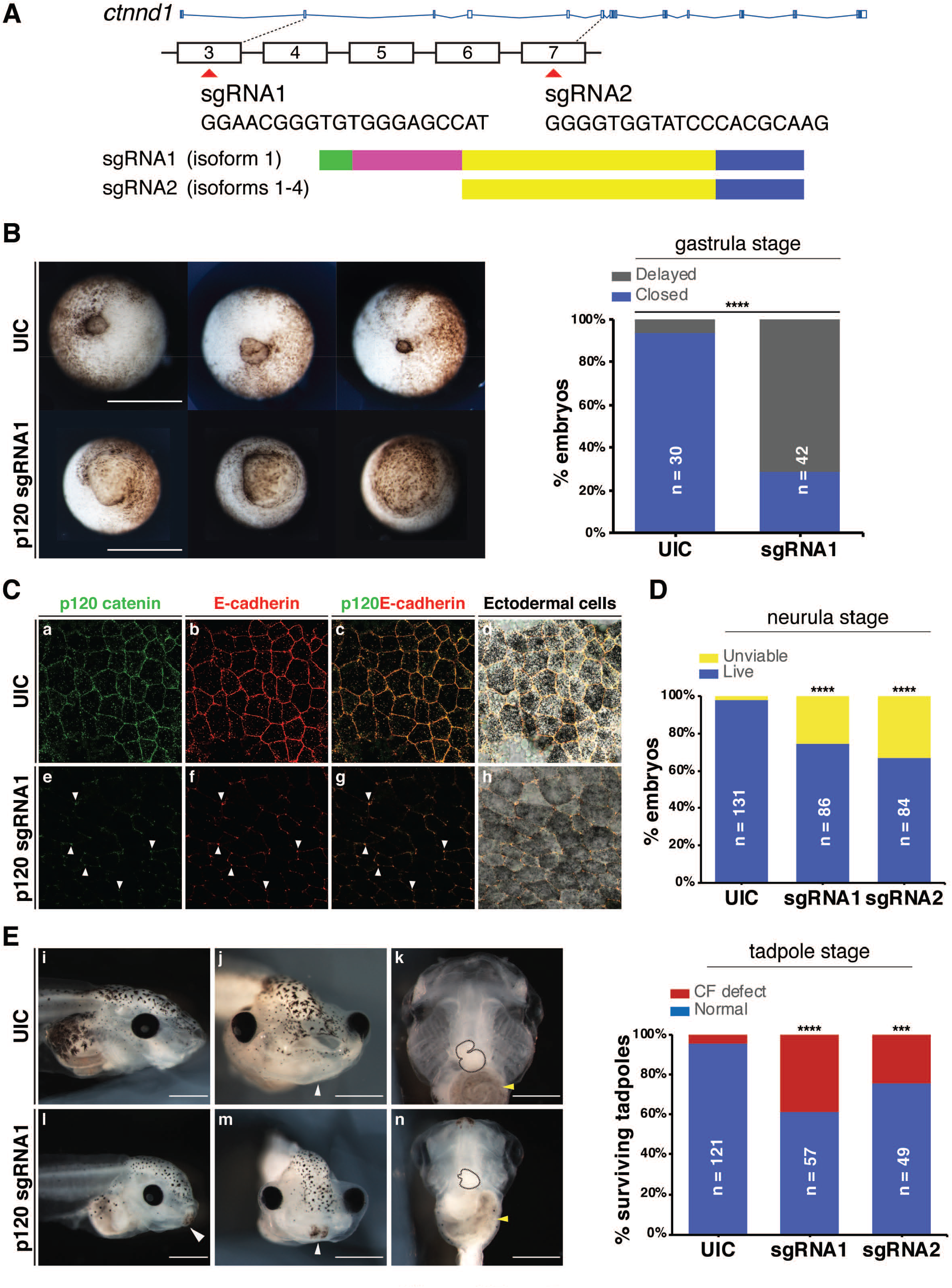
Ctnnd1 knockouts in Xenopus give rise to craniofacial and heart defects. [A] Embryos were injected at the one cell stage with single guide RNAs (sgRNA), sgRNA1 and sgRNA2 targeting exons 3 and 7, respectively. [B] Ventral view showing blastopores at stage 11. Embryos injected with sgRNA1 had delayed blastopore closure (bottom row) compared to un-injected controls (UIC) (top row). The bar chart shows quantitation. Scale bars = 100µm. [C] Confocal sections through the apical surface of ectodermal cells at stage 11 of embryos injected with sgRNA1 (e-h) and UICs (a-d). [C] (a-d) p120-catenin (a, green) is expressed in puncta at the cell membranes. E-cadherin (b, red) is expressed more evenly through the cell membranes. Both are colocalized at the cell-cell interface (c, d). Endogenous levels of p120-catenin and E-cadherin are diminished at the cell-cell interface in the sgRNA1-injected embryos (e-f). Residual p120-catenin and E-cadherin are seen in a spot-like pattern, only at the tricellular junctions (e-h, white arrowheads). [D] p120-catenin depletion led to lethality in embryos by the neurula stage. [E] Stage 46 tadpoles. [E] (i, l) Lateral views show a flattened profile in *p120* CRISPR tadpoles (l) compared to UICs (i). [E] (j, m) Frontal views showing a reduction in the size of mouth opening and a persistent cement gland (white arrowhead) in *p120* CRISPR tadpoles (m) compared to UICs (j). [E] (k,n) Ventral views showing a reduction in the size of craniofacial cartilages, altered cardiac looping (black-dashed outline) and altered gut coiling (yellow arrowhead) in *p120* CRISPR tadpoles (n) compared to UICs (k). Quantification of craniofacial defects in UIC and p120 depleted tadpoles. Scale bars = 100µm. sgRNA, single guide RNA; UIC, un-injected control; ****p<0.0001; ***p<0.001.

**Figure 8.**
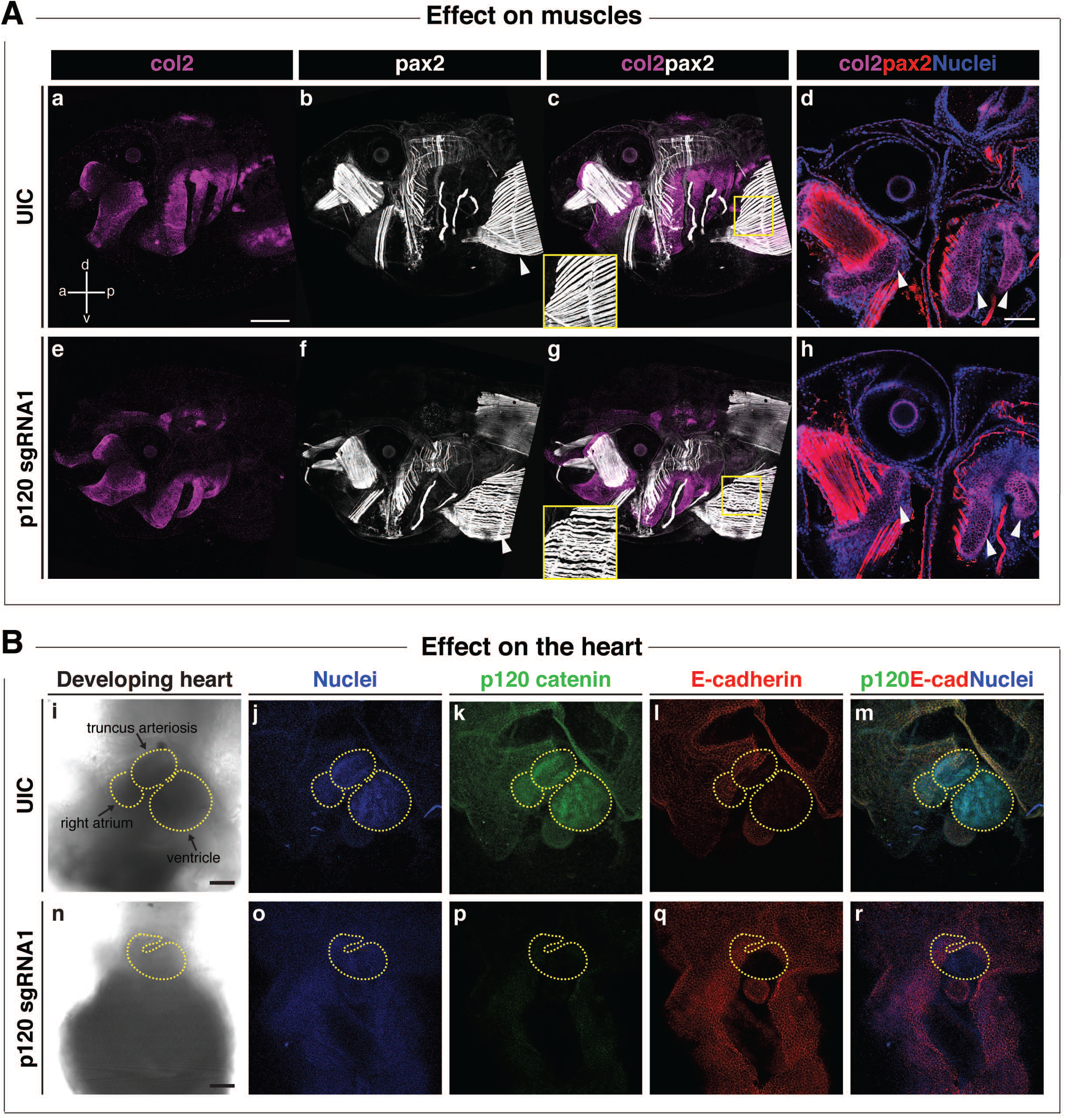
Ctnnd1 knockouts in Xenopus give rise to altered morphogenesis of the muscles and heart. [A] Immunofluorescent staining for collagen 2 (col2, magenta), muscle/pax2 (white) and nuclei (DAPI, blue); (a, anterior; p, posterior; d, dorsal; v, ventral). [A] (a, e) A lateral view of col2-positive branchial cartilages in UIC (a) and *p120* CRISPR mutant (e) reveals hypoplasia of mutant cartilages; however, cell morphology appears normal in *p120* CRISPR mutants (h) (d and h, white arrowheads). [A] (b-c, f-g) Pax2-expressing muscles revealed a defect in the fibril organization of the rectus abdominus muscle in the *p120* CRISPR tadpoles (f, white arrowhead) compared to the UIC muscles (b, white arrowhead); note insets in (c, g). [B] Ventral views of hearts of stage 46 tadpoles. Immunofluorescent staining for p120-catenin (green), E-cadherin (red) and DNA (blue). [B] (i-m) Controls; (n-r) *p120* CRISPR mutant tadpoles. Morphologic defects are evident in the size of the heart and directionality of the loops (compare control heart (i) to mutant heart (n), yellow-dashed outlines). [B] (k, p) p120-catenin is strongly expressed in the heart of UIC tadpoles (k) but is lost in *p120* CRISPR tadpoles (p). [B] (l, q) Note the absence of E-cadherin in the control and mutant hearts. Scale bars = 100µm.

**Figure 9.**
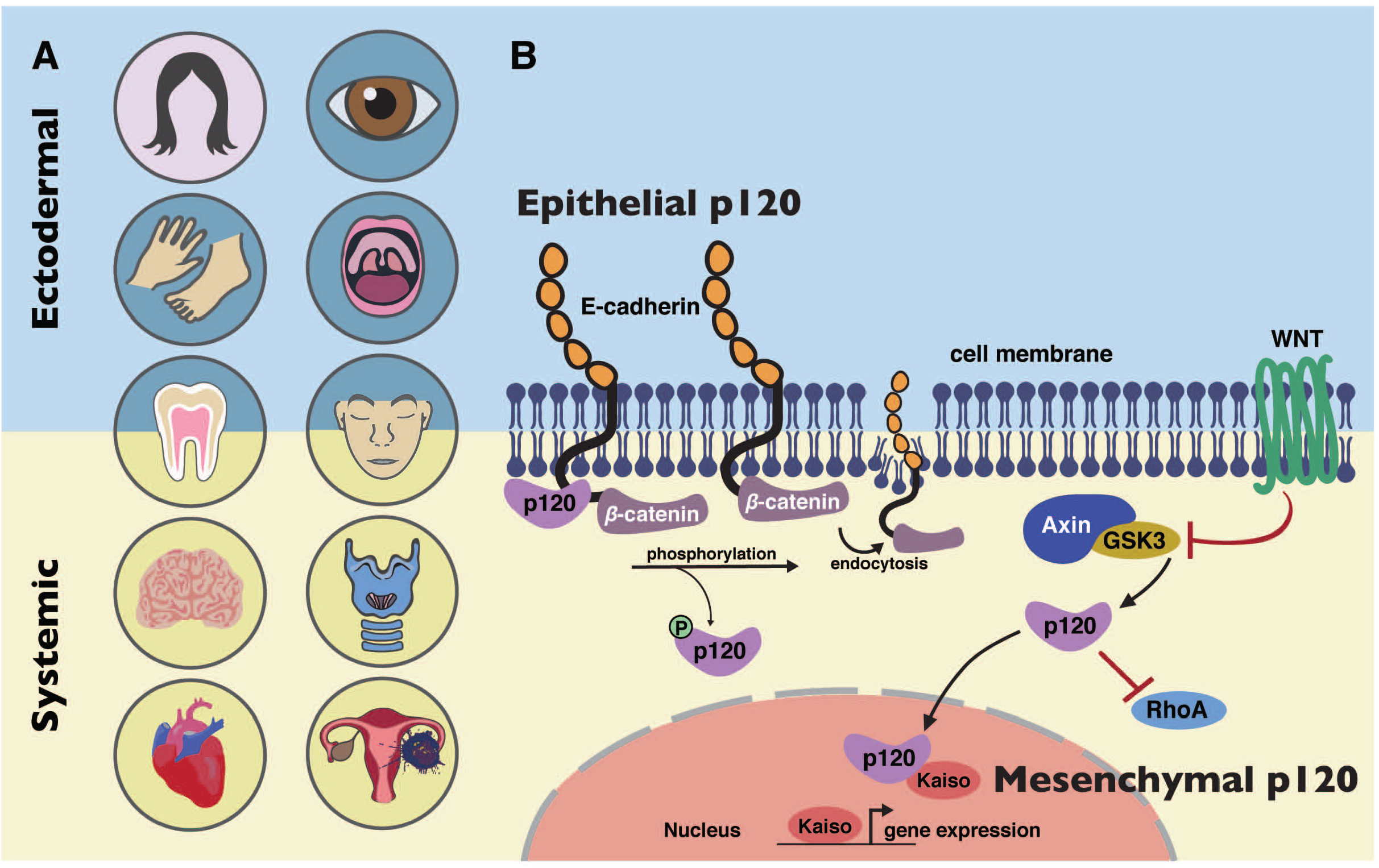
Model of CTNND1 function in systemic disease. [A] *CTNND1* mutations are not only implicated in conditions that affect epithelial structures but also systemic conditions that originate from mesenchymal roles of p120-catenin. Structures in pink circles have been described in previous publications on *CTNND1*^1, 2^; structures in blue circles have been implicated previously in *CTNND1*-related disorders^1, 2^ and in this study; structures in yellow circles have been identified in this study. [B] Blepharocheilodontic syndrome (BCD) is primarily due to disturbances in E-cadherin/p120 interactions. The inclusion of other organ systems described here highlights the involvement of other known molecular functions of p120, such as its role in the WNT signaling pathway and its interactions with Rho-GTPases, demonstrating its mesenchymal roles in producing these systemic conditions.

In addition to the phenotypes shared commonly across our cohort, some participants in this study had scoliosis, and one family reported two deceased children, who had bifid uvula, congenital cardiac disease (VSD, PDS), eye anomalies, developmental delay and chronic bowel immotility and gastroesophageal reflux disease; however, no genetic testing had been carried out. One patient presented at a young age with an ovarian dysgerminoma. To our knowledge, this is the first patient with a *CTNND1* variant associated with an early onset cancer, though p120 has been associated with cancer and tumorigenesis^23–25, 93, 94^. Finally, a number of patients reported in DECIPHER have copy number variants (CNV) affecting *CTNND1* (data not shown). Interestingly, both deletions and duplications have been associated with partially overlapping phenotypes. For instance, two patients with a deletion of less than 4MB had anomalies including bulbous nose, limb anomalies, delayed speech and language development, intellectual disability, nasal speech, ventricular septal defect, and cleft lip.

In summary, we demonstrate for the first time that p120 is not only involved in human conditions involving epithelial integrity, most likely caused by aberrant E-cadherin/p120 interactions, but also in other important intracellular functions (Figure 9). We conclude that *CTNND1-*related disorders span a spectrum of phenotypes ranging from multi-system involvement, to non-syndromic clefting. While further studies will be necessary to definitively understand the phenotype-genotype correlations, *CTNND1*, and perhaps *CDH1,* should be considered when patients present with characteristic craniofacial anomalies, congenital cardiac defects and neurodevelopmental disorders.

## Declarations

SAL is part owner of Qiyas Higher Health, a startup company unrelated to this work.

## Acknowledgements

We thank the patients and their families for their kind co-operation. We are grateful to the South Thames Cleft Team for their support, and to the Liu lab and colleagues in CCRB, especially Angela Gates, for support and feedback. The Human Developmental Biology Resource, which provided human samples, is funded by Joint MRC/Wellcome Trust (grant # 099175/Z/12/Z). This study makes use of **<u>DECIPHER</u>**, which is funded by the Wellcome Trust. The DDD study presents independent research commissioned by the Health Innovation Challenge Fund [grant number HICF-1009-003], a parallel funding partnership between Wellcome and the Department of Health, and the Wellcome Sanger Institute [grant number WT098051]. The views expressed in this publication are those of the author(s) and not necessarily those of Wellcome or the Department of Health. The research team acknowledges the support of the National Institute for Health Research, through the Comprehensive Clinical Research Network. *Xenopus* experiments were additionally supported by the European Xenopus Resource Centre, Portsmouth UK, the National Xenopus Resource USA and Xenbase. Work in the Liu lab is funded by BBSRC (KJL), British Heart Foundation (KJL/JG), the MRC (KJL), the Faculty of Dental Surgery Royal College of Surgeons of England - British Society of Paediatric Dentistry (FDS RCSEng-BSPD) Small Grant (RA/MTH/KJL), KSA (RA) and NIH/NIDDK DK099478 (DKM).

---

**Figure Supplemental 1. Clinical presentation of individuals with a CTNND1 mutation.**

[A] The eye phenotypes of the narrow palpebral fissures, the hooded eyelids and highly arched, thin lateral eyebrows were evident from a young age. [B] Ear anomalies included: low-set ears, sometimes asymmetric and/or small; overfolded helices of the external ears; a pre-auricular pit was also seen in one of the patients (data not shown). [C] Upper limb anomalies included: slightly shorter 5^th^ fingers as seen in Patients 3, 12 and 13; and a single transverse palmar crease on the right hand seen in both Patients 3 and 8. Lower limb anomalies included: 2,3-cutaneous syndactyly of the feet; sandal gaps and camptodactyly of the 2^nd^ toe as seen in Patients 12 and 13; a longer 4^th^ toe in Patient 6 and short toes in Patient 7.

**Figure Supplemental 2.**
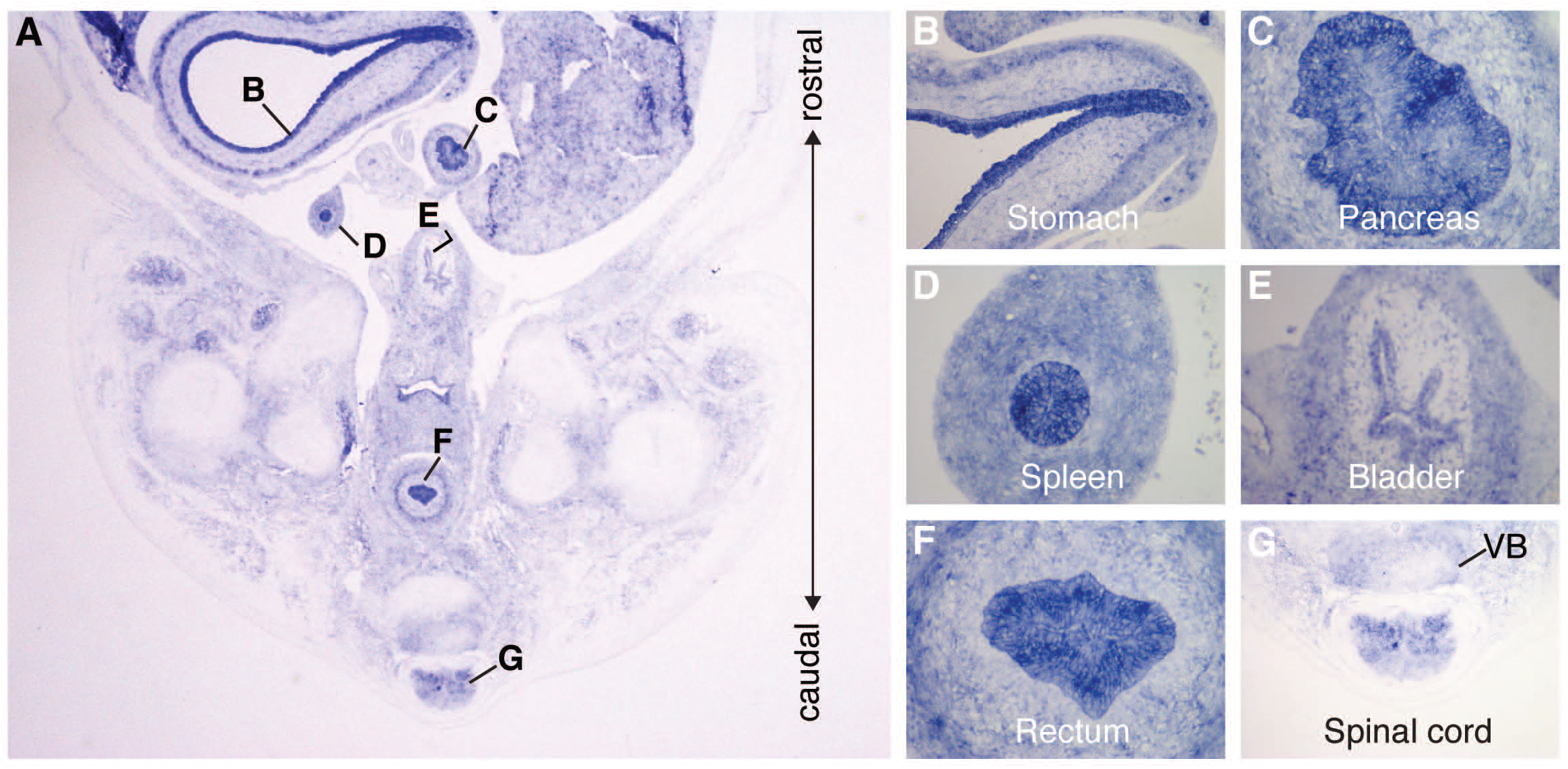
CTNND1 is expressed during relevant stages of human embryonic development. [A] Coronal cross-section through the torso at CS21. [B] *CTNND1* is expressed in the columnar epithelial lining of the stomach wall and continues through the pyloric part of the stomach. [C] Expression is seen in the islet of Langerhans in the pancreas. [D] Expression in the germinal center of the spleen. [E-G] Progressing caudally through the pelvis, *CTNND1* is expressed in the epithelial lining of the bladder [E], the rectum/hindgut [F], the spinal cord and vertebral body (VB) [G].

**Figure Supplemental 3.**
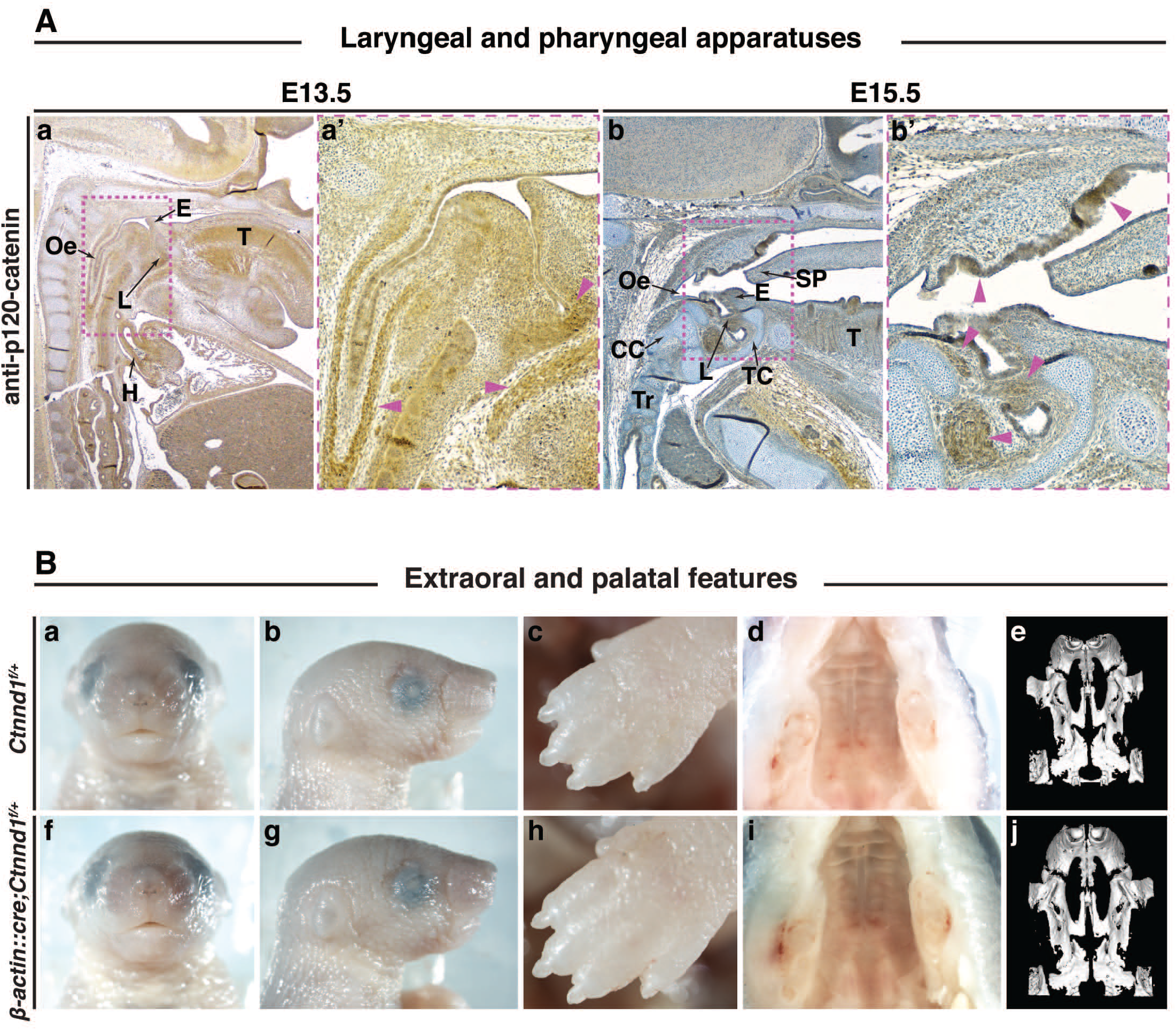
Mouse p120-catenin is expressed in the epithelial and mesenchymal compartments of the laryngeal and pharyngeal apparatuses. [A] Immunohistochemistry using the anti-phosphotyrosine p120-catenin antibody on sagittal sections through wild-type mice at E13.5 (a-a’) or E15.5 (b-b’). [a, b] Positive staining is seen in the epiglottis, esophagus and the larynx. [A] (a’, b’) Insets from (a and b, respectively). Muscles that express p120-catenin in the laryngeal and pharyngeal apparatuses are shown (pink arrowheads). Abbreviations: E, epiglottis; Oe, entrance to oesophagus; L, laryngeal auditus; H, heart; SP, soft palate; TC, thyroid cartilage; CC, cricoid cartilage; T, tongue; Tr, trachea **Heterozygosity in p120-catenin leads to normal facial and oral phenotypes.** [B] Shown are postnatal P2.5 mice. Heterozygous mutant *β-actin::cre/+;Ctnnd1^fl/+^* mice do not exhibit facial or lip anomalies (f-g) and are comparable to littermate controls (a-b). [B] (c, h) No limb anomalies are observed. [B] (d, i) Postnatal P1 mice. Intra-oral views of the palate of wild-type (d) and heterozygous mutant littermate (i), cleft palate defects were not observed. [B] (e, j) Microcomputed tomography (µCT) scans showed normal palates in P2.5 control (e) and heterozygous mutant littermate (j).

**Table S1:**
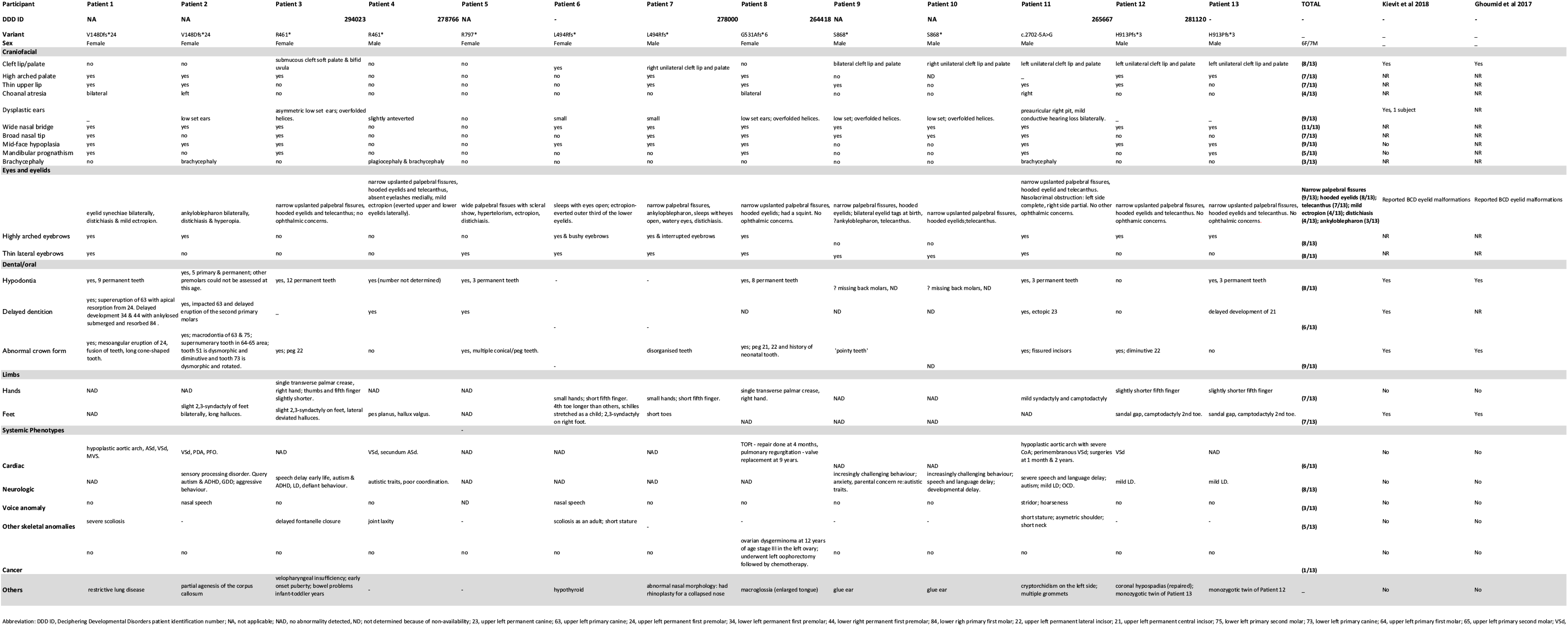
Clinical Details of Individuals with a *CTNND1* variant.

**Table S2:**
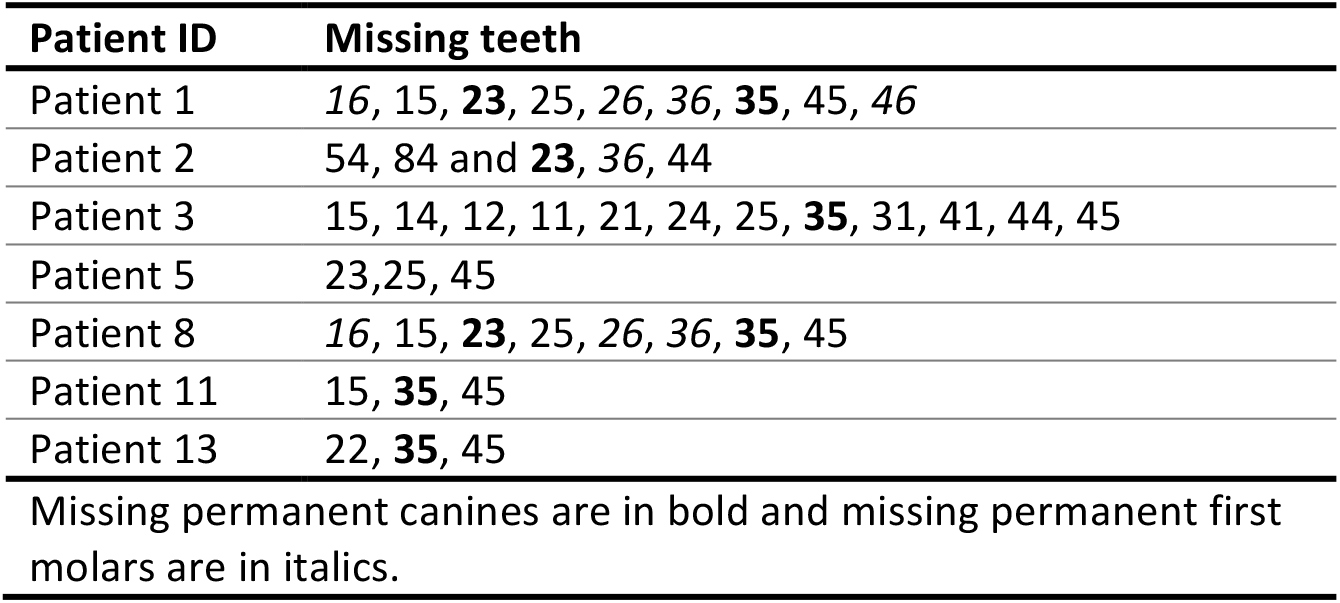
Reported congenitally missing teeth.

## REFERENCES

1. Kievit A, Tessadori F, Douben H, et al. Variants in members of the cadherin-catenin complex, CDH1 and CTNND1, cause blepharocheilodontic syndrome. Eur J Hum Genet. 2018;26(2):210–219.

2. Ghoumid J, Stichelbout M, Jourdain AS, et al. Blepharocheilodontic syndrome is a CDH1 pathway-related disorder due to mutations in CDH1 and CTNND1. Genetics in medicine: official journal of the American College of Medical Genetics. 2017;19(9):1013–1021.

3. Cox LL, Cox TC, Moreno Uribe LM, et al. Mutations in the Epithelial Cadherin-p120-Catenin Complex Cause Mendelian Non-Syndromic Cleft Lip with or without Cleft Palate. Am J Hum Genet. 2018;102(6):1143–1157.

4. Perez-Moreno M, Davis MA, Wong E, Pasolli HA, Reynolds AB, Fuchs E. p120-catenin mediates inflammatory responses in the skin. Cell. 2006;124(3):631–644.

5. Oas RG, Xiao K, Summers S, et al. p120-Catenin is required for mouse vascular development. Circ Res. 2010;106(5):941.

6. Marciano DK, Brakeman PR, Lee C-Z, et al. p120 catenin is required for normal renal tubulogenesis and glomerulogenesis. Development. 2011;138(10):2099–2109.

7. Hendley AM, Provost E, Bailey JM, et al. p120 Catenin is required for normal tubulogenesis but not epithelial integrity in developing mouse pancreas. Dev Biol. 2015;399(1):41–53.

8. Elia LP, Yamamoto M, Zang K, Reichardt LF. p120 catenin regulates dendritic spine and synapse development through Rho-family GTPases and cadherins. Neuron. 2006;51(1):43–56.

9. Davis MA, Reynolds AB. Blocked acinar development, E-cadherin reduction, and intraepithelial neoplasia upon ablation of p120-catenin in the mouse salivary gland. Developmental cell. 2006;10(1):21–31.

10. Bartlett JD, Dobeck JM, Tye CE, et al. Targeted p120-catenin ablation disrupts dental enamel development. PLoS One. 2010;5(9):e12703.

11. Ciesiolka M, Delvaeye M, Van Imschoot G, et al. p120 catenin is required for morphogenetic movements involved in the formation of the eyes and the craniofacial skeleton in Xenopus. J Cell Sci. 2004;117(18):4325–4339.

12. Geis K, Aberle H, Kühl M, Kemler R, Wedlich D. Expression of the Armadillo family member p120 cas 1B in Xenopus embryos affects head differentiation but not axis formation. Dev Genes Evol. 1998;207(7):471–481.

13. Ishiyama N, Lee S-H, Liu S, et al. Dynamic and static interactions between p120 catenin and E-cadherin regulate the stability of cell-cell adhesion. Cell. 2010;141(1):117–128.

14. Ireton RC, Davis MA, van Hengel J, et al. A novel role for p120 catenin in E-cadherin function. J Cell Biol. 2002;159(3):465–476.

15. Fukumoto Y, Shintani Y, Reynolds AB, Johnson KR, Wheelock MJ. The regulatory or phosphorylation domain of p120 catenin controls E-cadherin dynamics at the plasma membrane. Exp Cell Res. 2008;314(1):52–67.

16. Davis MA, Ireton RC, Reynolds AB. A core function for p120-catenin in cadherin turnover. J Cell Biol. 2003;163(3):525–534.

17. Reynolds AB, Daniel J, McCrea PD, Wheelock MJ, Wu J, Zhang Z. Identification of a new catenin: the tyrosine kinase substrate p120cas associates with E-cadherin complexes. Mol Cell Biol. 1994;14(12):8333–8342.

18. Anastasiadis PZ, Moon SY, Thoreson MA, et al. Inhibition of RhoA by p120 catenin. Nature cell biology. 2000;2(9):637.

19. Wildenberg GA, Dohn MR, Carnahan RH, et al. p120-catenin and p190RhoGAP regulate cell-cell adhesion by coordinating antagonism between Rac and Rho. Cell. 2006;127(5):1027–1039.

20. Park J-i, Kim SW, Lyons JP, et al. Kaiso/p120-catenin and TCF/β-catenin complexes coordinately regulate canonical Wnt gene targets. Developmental cell. 2005;8(6):843–854.

21. del Valle-Pérez B, Casagolda D, Lugilde E, et al. Wnt controls the transcriptional activity of Kaiso through CK1ε-dependent phosphorylation of p120-catenin. J Cell Sci. 2011;124(13):2298–2309.

22. Yanagisawa M, Anastasiadis PZ. p120 catenin is essential for mesenchymal cadherin– mediated regulation of cell motility and invasiveness. The Journal of cell biology. 2006;174(7):1087–1096.

23. Stairs DB, Bayne LJ, Rhoades B, et al. Deletion of p120-catenin results in a tumor microenvironment with inflammation and cancer that establishes it as a tumor suppressor gene. Cancer Cell. 2011;19(4):470–483.

24. Schackmann RC, Tenhagen M, van de Ven RA, Derksen PW. p120-catenin in cancer– mechanisms, models and opportunities for intervention. J Cell Sci. 2013;126(16):3515–3525.

25. Reynolds AB, Roczniak-Ferguson A. Emerging roles for p120-catenin in cell adhesion and cancer. Oncogene. 2004;23(48):7947.

26. Reynolds AB, Jenkins NA, Gilbert DJ, et al. The gene encoding p120cas, a novel catenin, localizes on human chromosome 11q11 (CTNND) and mouse chromosome 2 (Catns). Genomics. 1996;31(1):127–129.

27. Montonen O, Aho M, Uitto J, Aho S. Tissue distribution and cell type-specific expression of p120ctn isoforms. J Histochem Cytochem. 2001;49(12):1487–1495.

28. Aho S, Levänsuo L, Montonen O, Kari C, Rodeck U, Uitto J. Specific sequences in p120ctn determine subcellular distribution of its multiple isoforms involved in cellular adhesion of normal and malignant epithelial cells. J Cell Sci. 2002;115(7):1391–1402.

29. Hong JY, Oh I-H, McCrea PD. Phosphorylation and isoform use in p120-catenin during development and tumorigenesis. Biochimica et Biophysica Acta (BBA)-Molecular Cell Research. 2016;1863(1):102–114.

30. Keirsebilck A, Bonné S, Staes K, et al. Molecular cloning of the human p120ctnCatenin gene (CTNND1): Expression of multiple alternatively spliced isoforms. Genomics. 1998;50(2):129–146.

31. Mariner DJ, Wang J, Reynolds AB. ARVCF localizes to the nucleus and adherens junction and is mutually exclusive with p120 (ctn) in E-cadherin complexes. J Cell Sci. 2000;113(8):1481–1490.

32. Hatzfeld M. The p120 family of cell adhesion molecules. Eur J Cell Biol. 2005;84(2-3):205–214.

33. Gu D, Sater AK, Ji H, et al. Xenopus δ-catenin is essential in early embryogenesis and is functionally linked to cadherins and small GTPases. J Cell Sci. 2009;122(22):4049–4061.

34. Turner TN, Sharma K, Oh EC, et al. Loss of delta-catenin function in severe autism. Nature. 2015;520(7545):51–56.

35. Lu Q, Aguilar BJ, Li M, Jiang Y, Chen Y-H. Genetic alterations of δ-catenin/NPRAP/Neurojungin (CTNND2): functional implications in complex human diseases. Hum Genet. 2016;135(10):1107–1116.

36. Medina M, Marinescu RC, Overhauser J, Kosik KS. Hemizygosity of δ-catenin (CTNND2) is associated with severe mental retardation in cri-du-chat syndrome. Genomics. 2000;63(2):157–164.

37. Hofmeister W, Nilsson D, Topa A, et al. CTNND2-a candidate gene for reading problems and mild intellectual disability. J Med Genet. 2015;52(2):111–122.

38. Nivard M, Mbarek H, Hottenga J, et al. Further confirmation of the association between anxiety and CTNND2: replication in humans. *Genes*, Brain and Behavior. 2014;13(2):195–201.

39. Belcaro C, Dipresa S, Morini G, Pecile V, Skabar A, Fabretto A. CTNND2 deletion and intellectual disability. Gene. 2015;565(1):146–149.

40. Sirotkin H, O’Donnell H, DasGupta R, et al. Identification of a new human catenin gene family member (ARVCF) from the region deleted in velo-cardio-facial syndrome. Genomics. 1997;41(1):75–83.

41. Butts SC. The facial phenotype of the velo-cardio-facial syndrome. Int J Pediatr Otorhinolaryngol. 2009;73(3):343–350.

42. Shprintzen R, Goldberg R, Lewin M, et al. A new syndrome involving cleft palate, cardiac anomalies, typical facies, and learning disabilities: velo-cardio-facial syndrome. The Cleft palate journal. 1978;15(1):56–62.

43. Cho K, Lee M, Gu D, et al. Kazrin, and its binding partners ARVCF- and delta-catenin, are required for Xenopus laevis craniofacial development. Dev Dyn. 2011;240(12):2601–2612.

44. Deciphering Developmental Disorders S, McRae JF, Clayton S, et al. Prevalence and architecture of de novo mutations in developmental disorders. Nature. 2017;542:433.

45. Kaufman MH, Kaufman MH. The atlas of mouse development. Vol 428: Academic press London; 1992.

46. Lewandoski M, Meyers E, Martin G. Analysis of Fgf8 gene function in vertebrate development. Paper presented at: Cold Spring Harbor symposia on quantitative biology1997.

47. Lewis AE, Vasudevan HN, O’Neill AK, Soriano P, Bush JO. The widely used Wnt1-Cre transgene causes developmental phenotypes by ectopic activation of Wnt signaling. Dev Biol. 2013;379(2):229–234.

48. Khokha MK, Chung C, Bustamante EL, et al. Techniques and probes for the study of Xenopus tropicalis development. Dev Dyn. 2002;225(4):499–510.

49. Rual JF, Hirozane-Kishikawa T, Hao T, et al. Human ORFeome version 1.1: a platform for reverse proteomics. Genome Res. 2004;14(10b):2128–2135.

50. Wilkinson DG, Bailes JA, McMahon AP. Expression of the proto-oncogene int-1 is restricted to specific neural cells in the developing mouse embryo. Cell. 1987;50(1):79–88.

51. Karczewski KJ, Francioli LC, Tiao G, et al. Variation across 141,456 human exomes and genomes reveals the spectrum of loss-of-function intolerance across human protein-coding genes. bioRxiv. 2019:531210.

52. Xia X, Mariner DJ, Reynolds AB. Adhesion-associated and PKC-modulated changes in serine/threonine phosphorylation of p120-catenin. Biochemistry (Mosc). 2003;42(30):9195–9204.

53. Vinyoles M, Del Valle-Pérez B, Curto J, et al. Multivesicular GSK3 sequestration upon Wnt signaling is controlled by p120-catenin/cadherin interaction with LRP5/6. Mol Cell. 2014;53(3):444–457.

54. Sun D, Mcalmon KR, Davies JA, Bernfield M, Hay ED. Simultaneous loss of expression of syndecan-1 and E-cadherin in the embryonic palate during epithelial-mesenchymal transformation. Int J Dev Biol. 2003;42(5):733–736.

55. Mariner DJ, Davis MA, Reynolds AB. EGFR signaling to p120-catenin through phosphorylation at Y228. J Cell Sci. 2004;117(8):1339–1350.

56. Leopold C, De Barros A, Cellier C, Drouin-Garraud V, Dehesdin D, Marie J-P. Laryngeal abnormalities are frequent in the 22q11 deletion syndrome. Int J Pediatr Otorhinolaryngol. 2012;76(1):36–40.

57. Miyamoto RC, Cotton RT, Rope AF, et al. Association of anterior glottic webs with velocardiofacial syndrome (chromosome 22q11. 2 deletion). Otolaryngology—Head and Neck Surgery. 2004;130(4):415–417.

58. Fokstuen S, Bottani A, Medeiros PF, Antonarakis SE, Stoll C, Schinzel A. Laryngeal atresia type III (glottic web) with 22q11. 2 microdeletion: report of three patients. Am J Med Genet. 1997;70(2):130–133.

59. Shawlot W, Deng JM, Fohn LE, Behringer RR. Restricted β-galactosidase expression of a hygromycin-lacZ gene targeted to the β-actin locus and embryonic lethality of β-actin mutant mice. Transgenic Res. 1998;7(2):95–103.

60. Elder PK, French CL, Subramaniam M, Schmidt LJ, Getz MJ. Evidence that the functional beta-actin gene is single copy in most mice and is associated with 5’sequences capable of conferring serum-and cycloheximide-dependent regulation. Mol Cell Biol. 1988;8(1):480–485.

61. Tabler JM, Rigney MM, Berman GJ, et al. Cilia-mediated Hedgehog signaling controls form and function in the mammalian larynx. Elife. 2017;6.

62. Danielian PS, Muccino D, Rowitch DH, Michael SK, McMahon AP. Modification of gene activity in mouse embryos in utero by a tamoxifen-inducible form of Cre recombinase. Curr Biol. 1998;8(24):1323–S1322.

63. Paulson AF, Fang X, Ji H, Reynolds AB, McCrea PD. Misexpression of the Catenin p120ctn1A PerturbsXenopusGastrulation But Does Not Elicit Wnt-Directed Axis Specification. Dev Biol. 1999;207(2):350–363.

64. Bhattacharya D, Marfo CA, Li D, Lane M, Khokha MK. CRISPR/Cas9: an inexpensive, efficient loss of function tool to screen human disease genes in Xenopus. Dev Biol. 2015;408(2):196–204.

65. Corsten-Janssen N, Saitta SC, Hoefsloot LH, et al. More Clinical Overlap between 22q11.2 Deletion Syndrome and CHARGE Syndrome than Often Anticipated. Mol Syndromol. 2013;4(5):235–245.

66. Vissers LE, van Ravenswaaij CM, Admiraal R, et al. Mutations in a new member of the chromodomain gene family cause CHARGE syndrome. Nat Genet. 2004;36(9):955–957.

67. Wong MT, Schölvinck EH, Lambeck AJ, van Ravenswaaij-Arts CM. CHARGE syndrome: a review of the immunological aspects. Eur J Hum Genet. 2015;23(11):1451.

68. Goos JAC, Swagemakers SMA, Twigg SRF, et al. Identification of causative variants in TXNL4A in Burn-McKeown syndrome and isolated choanal atresia. Eur J Hum Genet. 2017;25(10):1126–1133.

69. Lopes VLGdS, Guion-Almeida ML, Rodini ESdO. Blepharocheilodontic (BCD) syndrome: expanding the phenotype? American Journal of Medical Genetics Part A. 2003;121(3):266–270.

70. Ababneh FK, Al-Swaid A, Elhag A, Youssef T, Alsaif S. Blepharo-cheilo-dontic (BCD) syndrome: Expanding the phenotype, case report and review of literature. American Journal of Medical Genetics Part A. 2014;164(6):1525–1529.

71. Hammond NL, Dixon J, Dixon MJ. Periderm: Life-cycle and function during orofacial and epidermal development. Paper presented at: Seminars in cell & developmental biology2017.

72. Abe R, Endo T, Shimooka S. Maxillary first molar agenesis and other dental anomalies. The Angle Orthodontist. 2010;80(6):1002–1009.

73. Nishi E, Masuda K, Arakawa M, et al. Exome sequencing-based identification of mutations in non-syndromic genes among individuals with apparently syndromic features. American journal of medical genetics Part A. 2016;170(11):2889–2894.

74. Lohi M, Tucker AS, Sharpe PT. Expression of Axin2 indicates a role for canonical Wnt signaling in development of the crown and root during pre- and postnatal tooth development. Dev Dyn. 2010;239(1):160–167.

75. Laurikkala J, Mikkola M, Mustonen T, et al. TNF signaling via the ligand–receptor pair ectodysplasin and edar controls the function of epithelial signaling centers and is regulated by Wnt and activin during tooth organogenesis. Dev Biol. 2001;229(2):443–455.

76. Wang B, Li H, Liu Y, et al. Expression patterns of WNT/β-CATENIN signaling molecules during human tooth development. Journal of molecular histology. 2014;45(5):487–496.

77. Liu F, Chu EY, Watt B, et al. Wnt/beta-catenin signaling directs multiple stages of tooth morphogenesis. Dev Biol. 2008;313(1):210–224.

78. Lammi L, Arte S, Somer M, et al. Mutations in AXIN2 cause familial tooth agenesis and predispose to colorectal cancer. The American Journal of Human Genetics. 2004;74(5):1043–1050.

79. Callahan N, Modesto A, Meira R, Seymen F, Patir A, Vieira A. Axis inhibition protein 2 (AXIN2) polymorphisms and tooth agenesis. Arch Oral Biol. 2009;54(1):45–49.

80. Mostowska A, Biedziak B, Jagodzinski PP. Axis inhibition protein 2 (AXIN2) polymorphisms may be a risk factor for selective tooth agenesis. J Hum Genet. 2006;51(3):262.

81. Song S, Zhao R, He H, Zhang J, Feng H, Lin L. WNT10A variants are associated with non-syndromic tooth agenesis in the general population. Hum Genet. 2014;133(1):117–124.

82. Mues G, Bonds J, Xiang L, et al. The WNT10A gene in ectodermal dysplasias and selective tooth agenesis. American Journal of Medical Genetics Part A. 2014;164(10):2455–2460.

83. van den Boogaard M-J, Créton M, Bronkhorst Y, et al. Mutations in WNT10A are present in more than half of isolated hypodontia cases. J Med Genet. 2012;49(5):327–331.

84. Mostowska A, Biedziak B, Zadurska M, Dunin-Wilczynska I, Lianeri M, Jagodzinski P. Nucleotide variants of genes encoding components of the Wnt signalling pathway and the risk of non-syndromic tooth agenesis. Clin Genet. 2013;84(5):429–440.

85. Lungova V, Verheyden JM, Sun X, Thibeault SL. β-Catenin signaling is essential for mammalian larynx recanalization and the establishment of vocal fold progenitor cells. Development. 2018;145(4):dev157677.

86. Park J-i, Ji H, Jun S, et al. Frodo links Dishevelled to the p120-catenin/Kaiso pathway: distinct catenin subfamilies promote Wnt signals. Developmental cell. 2006;11(5):683–695.

87. Kim SW, Park J-I, Spring CM, et al. Non-canonical Wnt signals are modulated by the Kaiso transcriptional repressor and p120-catenin. Nature cell biology. 2004;6(12):1212.

88. Buckingham M, Meilhac S, Zaffran S. Building the mammalian heart from two sources of myocardial cells. Nature Reviews Genetics. 2005;6(11):826.

89. Srivastava D, Thomas T, Lin Q, Kirby ML, Brown D, Olson EN. Regulation of cardiac mesodermal and neural crest development by the bHLH transcription factor, dHAND. Nat Genet. 1997;16(2):154.

90. Kochilas L, Merscher-Gomez S, Lu MM, et al. The role of neural crest during cardiac development in a mouse model of DiGeorge syndrome. Dev Biol. 2002;251(1):157–166.

91. Eley L, Alqahtani AM, MacGrogan D, et al. A novel source of arterial valve cells linked to bicuspid aortic valve without raphe in mice. Elife. 2018;7.

92. Peterson JC, Chughtai M, Wisse LJ, et al. Nos3 mutation leads to abnormal neural crest cell and second heart field lineage patterning in bicuspid aortic valve formation. Disease Models & Mechanisms. 2018:dmm. 034637.

93. Smalley-Freed WG, Efimov A, Burnett PE, et al. p120-catenin is essential for maintenance of barrier function and intestinal homeostasis in mice. The Journal of clinical investigation. 2010;120(6):1824–1835.

94. Lehman HL, Yang X, Welsh PA, Stairs DB. p120-catenin down-regulation and epidermal growth factor receptor overexpression results in a transformed epithelium that mimics esophageal squamous cell carcinoma. The American journal of pathology. 2015;185(1):240–251.

